# Cueing motor memory reactivation during NREM sleep engenders learning-related changes in precuneus and sensorimotor structures

**DOI:** 10.1101/2022.01.27.477838

**Authors:** Martyna Rakowska, Paulina Bagrowska, Alberto Lazari, Miguel Navarrete, Mahmoud E. A. Abdellahi, Heidi Johansen-Berg, Penelope A. Lewis

**Affiliations:** Cardiff University Brain Research Imaging Centre (CUBRIC), School of Psychology, Cardiff University, Cardiff, UK; Experimental Psychopathology Lab, Institute of Psychology, Polish Academy of Sciences, Warsaw, Poland; Wellcome Centre for Integrative Neuroimaging, FMRIB, Nuffield Department of Clinical Neurosciences, University of Oxford, Oxford, UK; Department of Biomedical Engineering, Universidad de los Andes, Bogotá, Colombia

**Author notes:** Corresponding author. Cardiff University Brain Research Imaging Centre (CUBRIC), School of Psychology, Cardiff University, Maindy Rd, Cardiff, CF24 4HQ, UK. Email address (M. Rakowska).

## Abstract

Memory reactivation during Non-Rapid Eye Movement (NREM) sleep is important for memory consolidation but it remains unclear exactly how such activity promotes the development of a stable memory representation. We used Targeted Memory Reactivation (TMR) in combination with longitudinal structural and functional MRI to track the evolution of a motor memory trace over 20 days. We show that repeated reactivation of motor memory during sleep leads to increased precuneus activation 24 h post-TMR. Interestingly, a decrease in precuneus activity over the next 10 days predicts longer-term cueing benefit. We also find both functional and structural changes in sensorimotor cortex in association with effects of TMR 20 days post-encoding. These findings demonstrate that TMR can engage precuneus in the short-term while also impacting on task-related structure and function over longer timescales.

## 1 INTRODUCTION

Memory consolidation is a process through which newly encoded memories become more stable and long-lasting. Consolidation is thought to involve repeated reinstatement, or reactivation of memory traces which allows their re-coding from short-term to long-term store^1^. Indeed, reactivation of learning-related brain activity patterns during sleep has been shown to predict subsequent memory performance^2,3^ and thus to play a critical role in memory consolidation^4,5^. However, it is unclear exactly how such offline rehearsal promotes the development of a stable memory representation. Here, we set out to investigate the neural plasticity underlying memory reactivation during sleep using Targeted Memory Reactivation (TMR) and magnetic resonance imaging (MRI).

TMR has recently emerged as a tool to study the mechanisms of memory reactivation. The technique involves re-presenting learning-associated cues during sleep^6^, thereby triggering reactivation of the associated memory representation and biasing their consolidation^7^. In humans, this manipulation shows strong behavioural effects (e.g., procedural memories TMR ^8–11^), resulting in better recall of memories that were cued through TMR compared to those that were not cued. Functional activity associated with cueing has been investigated during and immediately after sleep^6,10,12,13^. However, little is known about precisely how the memory representations targeted by TMR evolve over longer time periods. We have previously reported behavioural effects of procedural memory cueing during sleep twenty days post-manipulation^11^. Yet, the functional plasticity underlying such benefits is unknown. It also remains to be established whether TMR can impact on brain structure and which regions support sleep-dependent memory consolidation in the long term.

In this study, we used TMR to determine if repeated reactivation of a motor memory trace during sleep engenders learning-related changes in the brain. We tracked such impacts over several weeks using functional and structural brain imaging (Fig.1A) and hypothesized that memory cueing during sleep would lead to rapid plasticity within the precuneus, a structure which houses newly formed memory representations or ‘engrams’^14^. This region was of special interest since it has been shown to respond to repeated learning-retrieval epochs which help to strengthen a memory^14^ and can be thought of as a proxy for memory reactivation in sleep^15^. However, we expected the long-term storage of the memory engram to prevail in the task-related areas. We chose to focus specifically on a procedural memory task because the importance of sleep in procedural memory consolidation is well established^16,17^. Furthermore procedural memory task improvements^18^ and the associated structural changes^19^ have been shown to persist over time, with the same being true for the TMR effects^11^. Our participants were trained on two motor sequences in a Serial Reaction Time Task (SRTT). Each of these was associated with a different set of auditory tones (Fig.1B) but only one was reactivated during subsequent NREM sleep (Fig.1C). During learning and two post-sleep re-test sessions (24 h and 10 days post-TMR), participants were scanned with structural MRI (T1-weighted, T1w) and functional MRI (fMRI), the latter being acquired during SRTT performance. We were thus able to compare brain activity during the cued and uncued sequence performance, as well brain structure, both after one night and after 10 days. Twenty days post-TMR participants were again re-tested on the SRTT, now outside the scanner, allowing examination of the long-term impacts of TMR on behaviour. The resultant dataset enabled us to investigate how the impact of cueing evolves across time, as well as study the relationships between structural, functional, and behavioural plasticity post-TMR.

**Fig. 1.**
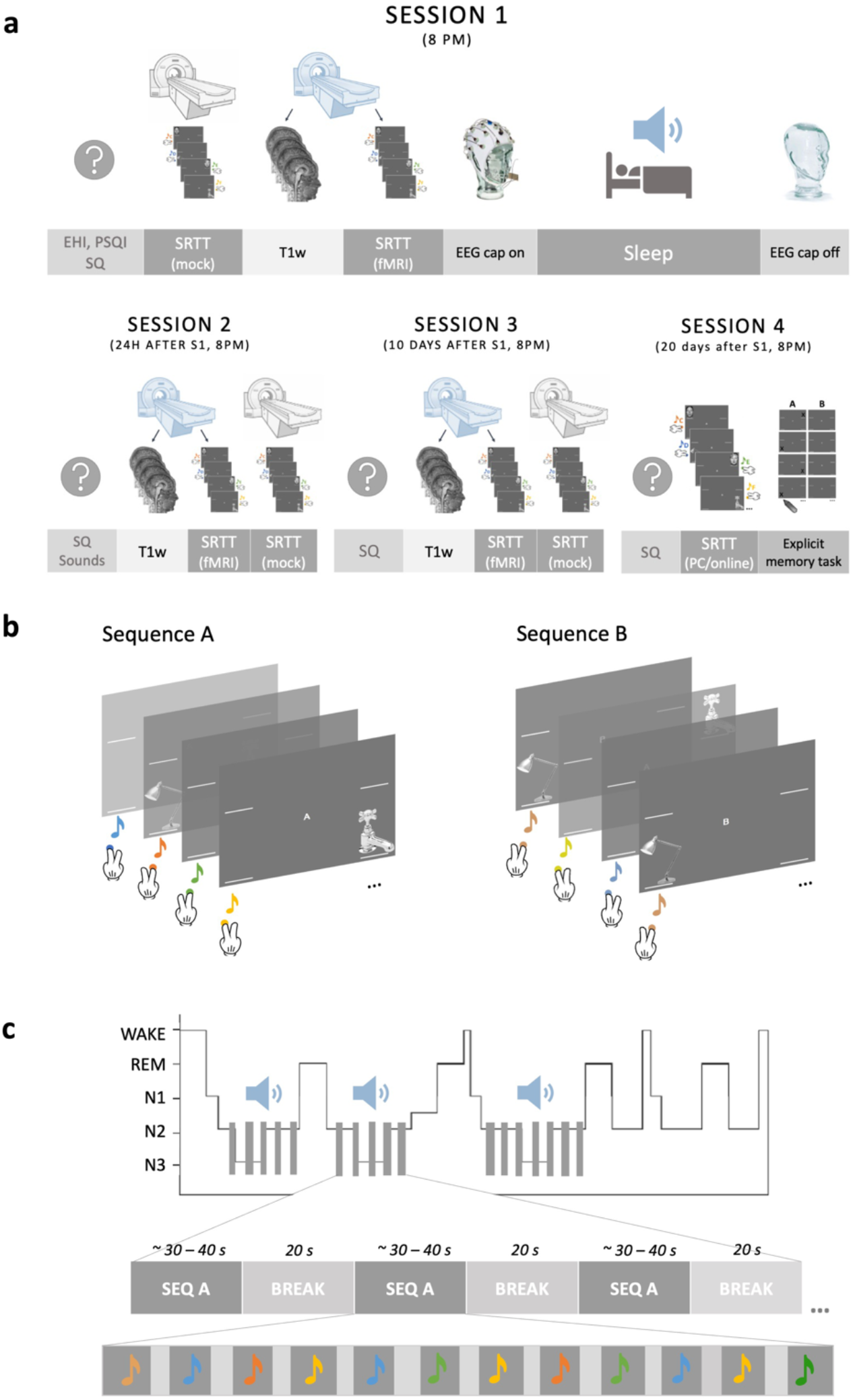
Study design and methods. **(a)** A schematic representation of the experimental sessions. SRTT and one or more questionnaires were delivered in each session. During S1-S3, SRTT was split in half, with the first half completed in the 0T ‘mock’ scanner (grey) and the second half in the 3T MRI scanner during fMRI acquisition (blue) (S1), or vice versa (S2-S3). T1w data was always acquired before fMRI. S1 also involved a stimulation night in the lab which the participants spent asleep and with the electroencephalography (EEG) cap on. During S4 SRTT data were acquired outside the MRI scanner and an explicit memory task was delivered at the very end of the study. **(b)** Two sequences of the SRTT. Only the first few trials are shown. Visual cues appeared at the same time as the auditory cues and the participants were instructed to push the key/button corresponding to the image location as quickly and accurately as possible. **(c)** TMR protocol. Tones associated with one sequence were played during stable N3 and N2 (grey bars on the hypnogram). One repetition of the cued sequence (dark grey rectangles) was followed by a 20 s break during which no sounds were played (light grey rectangles). Each sequence repetition comprised 12 tones (depicted as coloured notes) with inter-trial interval jittered between 2,500 and 3,500 ms (light grey vertical bars). S1-S4: Session 1 – Session 4; EHI: Edinburgh Handedness Inventory; PSQI: Pittsburgh Sleep Quality Index; SQ: Stanford Sleepiness Scale Questionnaire; SRTT: Serial Reaction Time Task; fMRI: functional Magnetic Resonance Imaging; T1w: T1-weighted scan.

## 2 RESULTS

### 2.1 SRTT

#### 2.1.1 REACTION TIME AND SEQUENCE SPECIFIC SKILL

Analysis of baseline SRTT performance indicated that participants learned both sequences before sleep and confirmed that any post-sleep differences between the sequences can be regarded as the effect of TMR (see *Supplementary Notes: Baseline SRTT performance* and Table S1). Fig.2A shows the mean reaction time (± SEM) for both hands (BH) trials of each SRTT block over the whole length of the study.

**Fig. 2.**
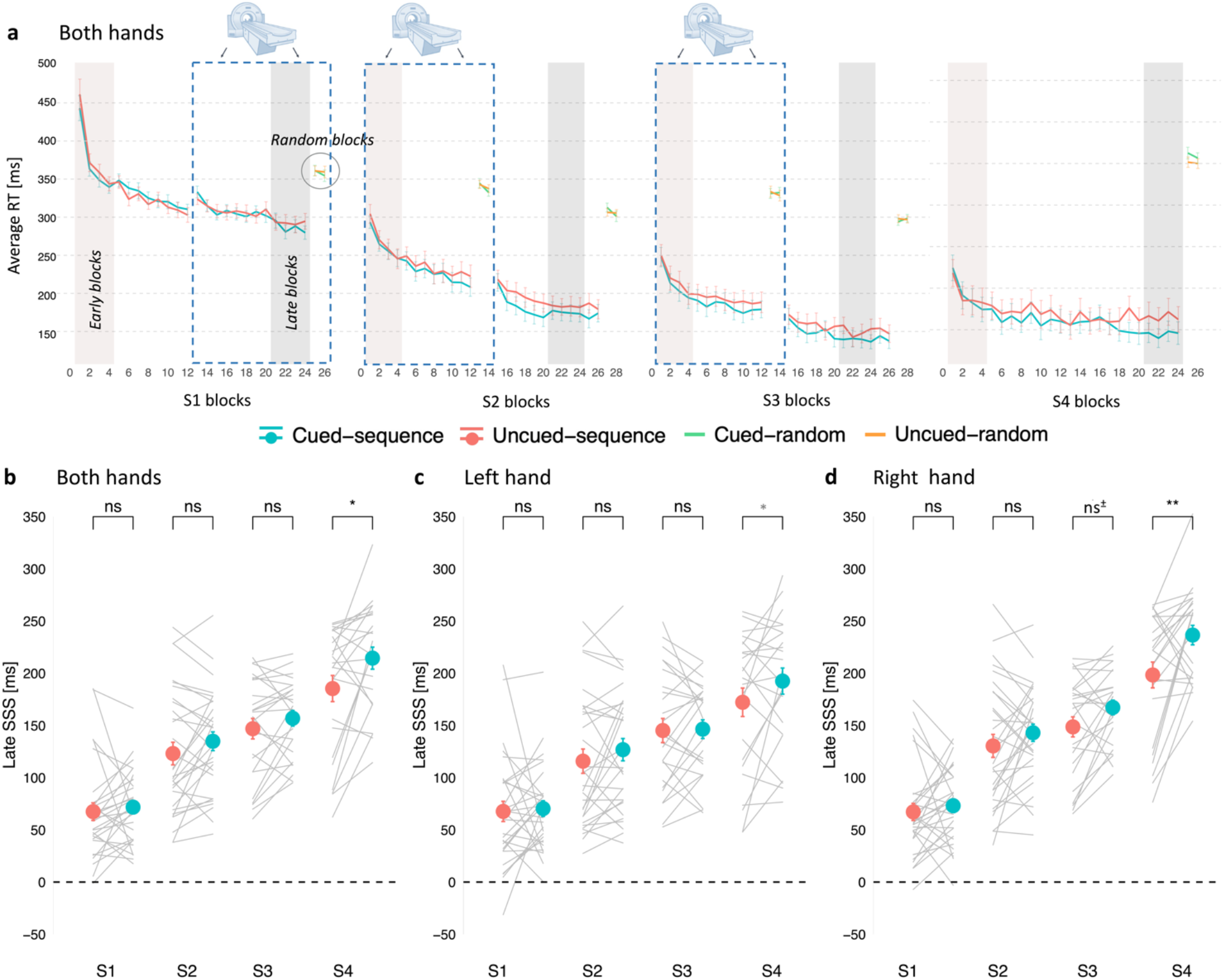
A single night of TMR benefits procedural memories up to 20 days later. **(a)** Mean reaction time for both hands trials of the cued sequence (blue), uncued sequence (red) and random blocks (green and orange) of the SRTT performed before sleep (S1), 24 h post-TMR (S2), 10 days post-TMR (S3) and 20 days post-TMR (S4). Error bars depict SEM. Vertical bars mark the first (brown) and last (grey) four sequence blocks used to determine early and late sequence specific skill (SSS), respectively. Blue dashed rectangle frames mark the SRTT blocks performed during fMRI acquisition. **(b-d)** Mean late SSS for the cued (blue dots) and uncued (red dots) sequence plotted against experimental sessions (S1-S4). Both hand trials **(b)**, left-hand trials **(c)** and right-hand trials **(d)** are shown separately. Error bars depict SEM. Grey lines represent individual participants. For (a-d): n = 30 for S1-S2, n = 25 for S3, n = 24 for S4. S1-4: Session 1-4; RT: reaction time; SSS: Sequence Specific Skill. **p < 0.001; *p < 0.05; ns^±^: non-significant trend (p = 0.060); ns: non-significant; p-values uncorrected; when adjusted for multiple comparisons using Holm’s correction the effect of TMR at S4 remained significant only for (B) and (D) (black *) but not for (C) (grey *).

Post-sleep SRTT re-test sessions occurred 24.67 h (SD: 0.70) (S2), 10.48 days (SD: 0.92) (S3), and 20.08 days (SD: 0.97) (S4) after session 1 (S1). SRTT performance was measured by subtracting the mean reaction time on the last or first four blocks of each sequence from that of the random blocks, thereby providing a measure of both late and early sequence specific skill (SSS). To test the effect of cueing on the SSS (either early or late) over time we fitted a linear mixed effects model to each dataset (both hands, BH; left hand, LH; right hand, RH) separately, with TMR and session entered as fixed effects, and participant entered as a random effect. Results of all the likelihood ratio tests comparing the full model against the model without the fixed effect of interest are shown in Table S2A-C.

The linear mixed effect analysis revealed a main effect of session on both early (X^2^(2) = 175.77, p < 0.001; Table S2Ai) and late SSS (X^2^(2) = 93.04, p < 0.001; Table S2Aii) for the BH dataset, with similar results for the LH (Table S2B) and RH datasets (Table S3C). Post-hoc comparisons showed a difference between subsequent sessions (S2 vs S3, S3 vs S4) (p_adj_ < 0.002; Table S3), suggesting continuous learning over time. All p_adj_ values are Holm-corrected.

Inclusion of TMR as a fixed effect in the BH dataset improved model fit across all sessions (S2-S4) for late SSS (X^2^(1) = 11.01, p = 0.001; Table S2Aii), but not early SSS (X^2^(1) = 1.55, p = 0.214; Table S2Ai). Similar results were revealed for LH (Table S2B) and RH datasets (Table S3C). Thus, the linear mixed effects analysis points to a main effect of TMR on the late SSS across all post-stimulation sessions. Given our previous findings on this task^11^, we expected better performance on the cued sequence at S3 (10 days post-stimulation) but wanted to determine if this TMR benefit (i.e., the difference between the SSS of the two sequences) persists until S4 (20 days post-stimulation). Likewise, we also aimed to probe if the TMR benefit emerges at S2 (24 h poststimulation) as other studies would suggest^10^, or whether it is non-significant at that time point (as in ^11^). Hence, we performed post-hoc comparisons to reveal the session(s) during which late SSS differed between the two sequences. Contrary to our expectations, we found a significant difference between the cued and uncued sequence performance at S4 (p_adj_ = 0.004) but not at S2 (p_adj_ = 0.282) or S3 (p_adj_ = 0.282) for the BH dataset (Table S4A, Fig.2B). Similar results were found for the RH dataset (S2: p_adj_ = 0.163; S3: p_adj_ = 0.119; S4: p_adj_ = 0.001; Table S4C, Fig.2D). However, for the LH dataset, the TMR benefit at S4 (p_uncorr_ = 0.040) did not survive Holm correction (S4: p_adj_ = 0.121; S2: p_adj_ = 0.421; S3: p_adj_ = 0.890) (Table S4B, Fig.2C). Together, these findings point to a main effect of TMR across all post-stimulation sessions, with the difference between the cued and uncued sequence being the strongest 20 days post-TMR, particularly for the dominant hand. Nevertheless, it is worth noting that although we do report a main effect of hand on the SSS (better performance for the dominant hand; p < 0.001), there was no interaction between hand and TMR (p > 0.05).

#### 2.1.2 CUEING BENEFIT ACROSS TIME

To explore how the TMR effect evolves over time, we calculated the difference between late SSS of the cued and uncued sequence for each session, which we refer to as the (late) cueing benefit. Next, we used a linear mixed effects analysis to determine if the cueing benefit changes across post-stimulation time. Inclusion of the number of days post-TMR as the fixed effect improved model fit on the extent of cueing benefit for BH (χ2(2) = 3.97, p = 0.046; Fig.3A) and RH dataset (χ2(2) = 6.58, p = 0.010; Fig.3C), but not for LH dataset (χ2(2) = 0.74, p = 0.391; Fig.3B) (Table S5). Interestingly, there was neither a main effect of hand nor an interaction between hand and session (p > 0.05; Table S5D). These results suggest that the effects of TMR develop in a gradual time-dependent manner.

**Fig. 3.**
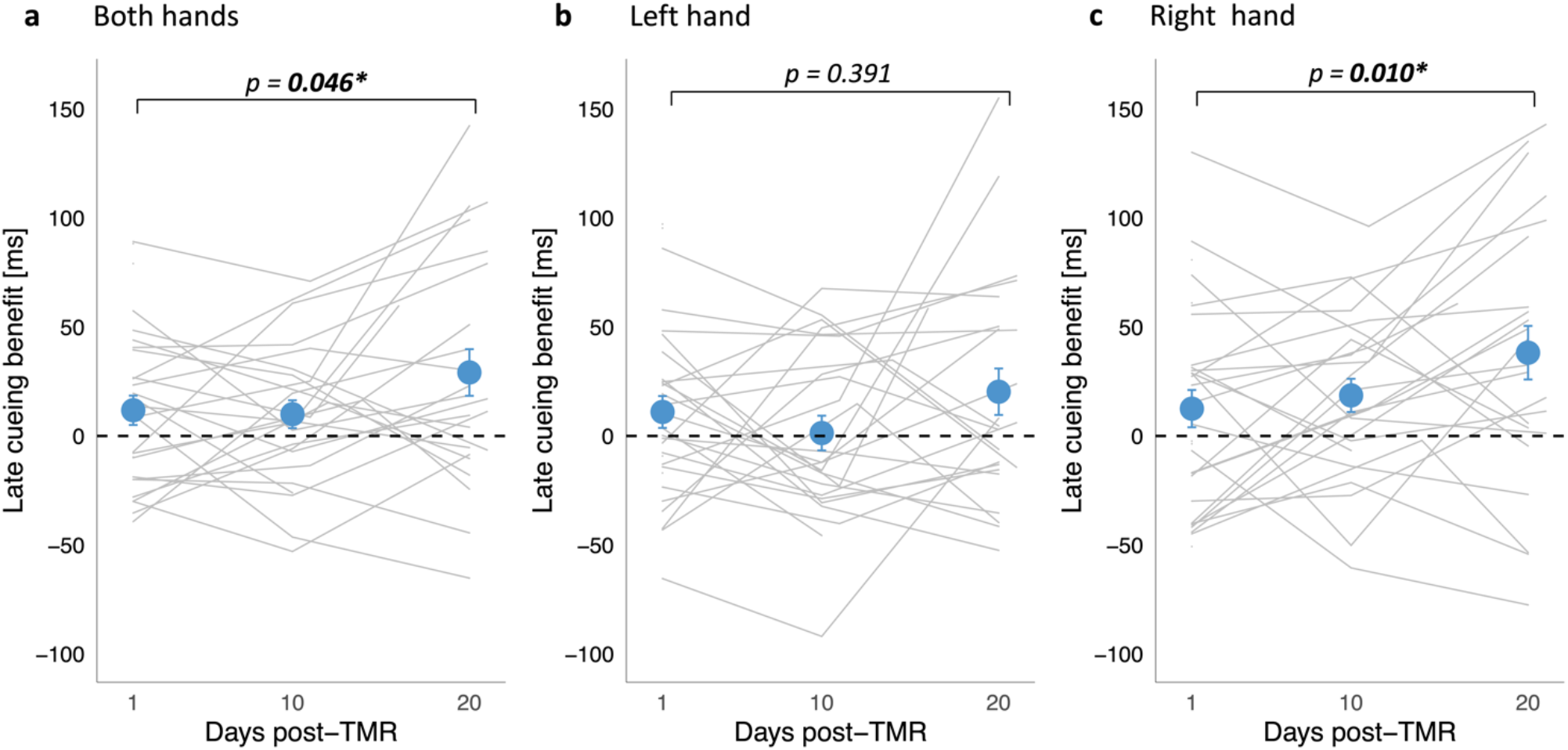
Cueing benefit over time. Mean late SSS on the uncued sequence subtracted from the cued sequence for both hands **(a),** left hand **(b)** and right hand **(c)**, plotted over time (number of days post-TMR). The effect of time was significant for both-hands dataset and right-hand datasets. Blue dots represent mean ±SEM calculated for S2, S3 and S4. Grey lines represent cueing benefit for each subject. N = 30 for S2, n = 25 for S3; n = 24 for S4. S2-S4: Session 2-4.

### 2.2 CORRELATIONS WITH SLEEP STAGES

To determine the relationship between sleep parameters derived from sleep stage scoring (Table S6) and the behavioural effect of our manipulation, we correlated the percentage of time spent in stage 2 (N2) and stage 3 (N3) of NREM sleep (the two target stages for our stimulation) with the cueing benefit at each session (S2, S3, S4) and for each dataset (BH, LH, RH) separately. Results are presented in Table S7, with no correlation surviving FDR correction (p_adj_ > 0.05).

### 2.3 SLEEP SPINDLES

Given the well-known involvement of sleep spindles in motor sequence memory consolidation^20^, we set out to describe electrophysiological changes within the spindle frequency in relation to the cueing procedure. The average spindle density over the task related regions was higher in N2 than in N3 during both the cue period (0-3.5 s after cue onset; t(28) = 4.48, p < 0.001) and the no-cue period (3.5-20 s after the onset of the last cue in the sequence; t(28) = 4.23, p < 0.0001) (paired-samples t-test). Next, we compared spindle density over left and right motor areas during N2 and N3 combined. The analysis revealed higher spindle density over the left versus right motor areas for the cue period (t(28) = 2.59, p = 0.015) but not for the no-cue period (t(28) = 1.98, p = 0.057) (paired-samples t-test). As in our previous study^11^, we also found that the average spindle density during the cue period was higher than during the no-cue period (t(28) = 4.37, p < 0.001; paired-samples t-test, Fig.4A-B), suggesting that cueing may elicit sleep spindles. Spindle density and the number of spindle events during each period and sleep stage are summarised in Table S8.

**Fig. 4.**
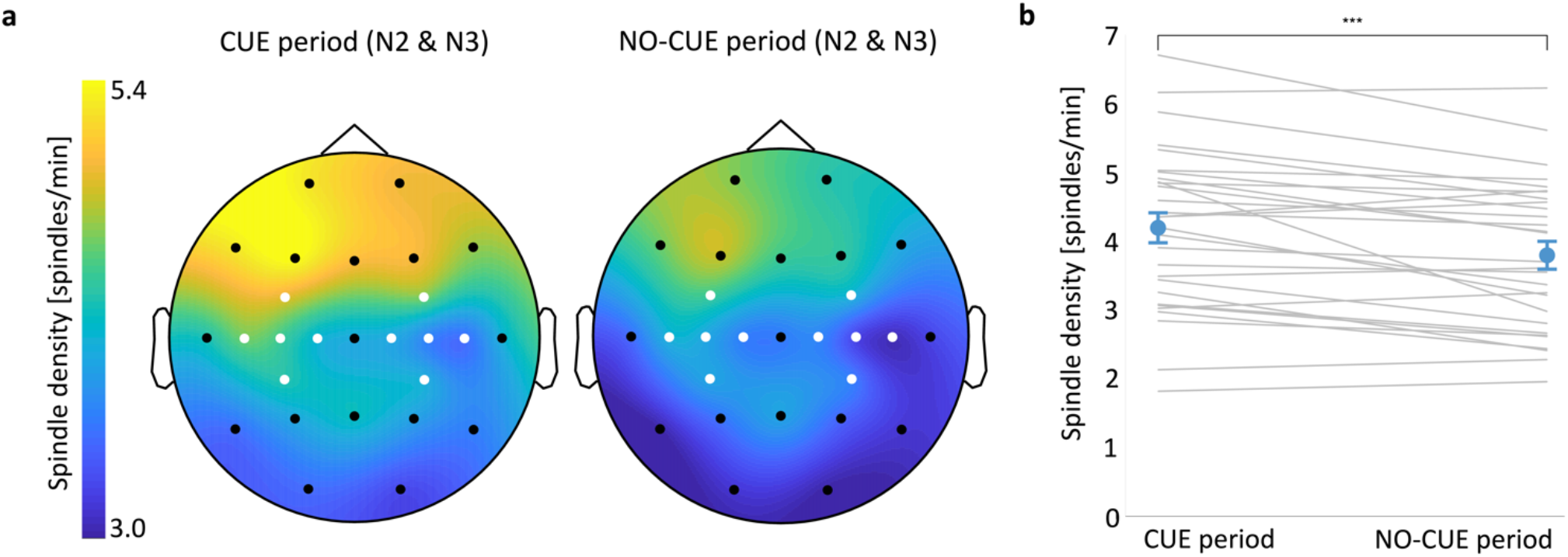
Spindle density increases immediately upon cue onset. **(a)** Topographic distribution of spindle density (spindles per min) in the cue (left) and no-cue (right) period of NREM sleep (N2 and N3 combined). Motor channels in white. **(b)** Spindle density averaged over motor channels during the cue period was higher than during the no-cue period. Blue dots represent mean ±SEM. Grey lines represent individual subjects. *** p = 0.001. N2-N3: stage 2 **–** stage 3 of NREM sleep. n = 29.

Spindle-related changes over brain regions involved in learning^21^ often predict behavioural performance^22^. However, we found no correlation between spindle density averaged over bilateral motor regions and cueing benefit for the BH dataset (p_adj_ > 0.05, Table S9).

### 2.4 TMR-RELATED CHANGES IN FMRI RESPONSE

We next examined BOLD responses to the cued and uncued sequence over time. To test our hypothesis that procedural memory cueing during sleep would engender learning-related changes within precuneus, we performed a TMR-by-Session ANOVA of the fMRI data acquired during sequence performance at S2 and S3. In line with our hypothesis, the analysis revealed increased activity in the right precuneus (8, −72, 58) for the main effect TMR (cued vs uncued sequence across S2 and S3) (peak F = 22.67, p = 0.032; Table S10A). We have previously shown cueing-related functional activity the morning after TMR^10^. Similarly, microstructural plasticity and functional engagement of posterior parietal cortex (PPC) has been detected relatively quickly after learning^14^. Thus, we expected functional activity changes at S2. Indeed, one-way t-tests on the [cued > uncued] contrast revealed increased activity in the dorsal-anterior subregion of left precuneus (−9, −62, 66) 24 h post-TMR (peak T = 4.79, p = 0.020; Fig.5A-B, Table S10B, Fig.S1A). Given our behavioural findings at S4, we also sought to determine if a TMR effect could be detected in fMRI data at S3, but no difference between cued and uncued activity was found at this time point (p > 0.05). These results show that TMR alters functional activity in precuneus, with the TMR-related increase in functional response apparent relatively quickly (i.e., 24 h) post-stimulation.

**Fig. 5.**
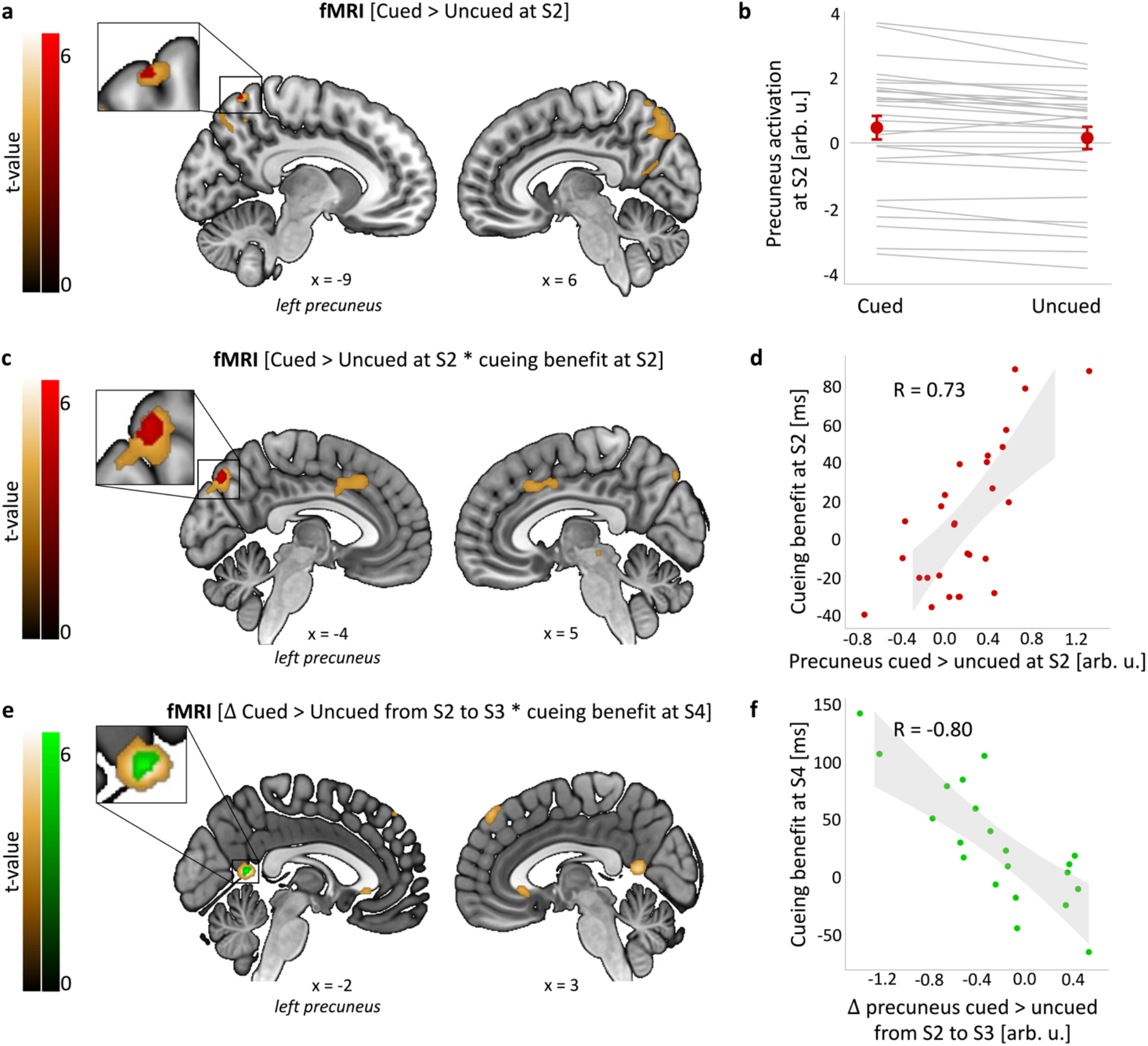
TMR-related functional activity in precuneus. **(a-b)** TMR-dependent increase in left precuneus activity 24 h post-stimulation. **(cd)** Activity for the [cued > uncued] contrast in the left precuneus at S2 is positively associated with behavioural cueing benefit at the same time point. **(e-f)** Change in activity from S2 to S3 for the [cued > uncued] contrast in the left precuneus is negatively associated with behavioural cueing benefit at S4. (**a, c, e**) Group level analysis. In red/green, colour-coded t-values for each contrast thresholded at a significance level of p_FWE_ < 0.05, corrected for multiple voxel-wise comparisons within a pre-defined ROI for bilateral precuneus. Increased activity shown in red; decreased activity shown in green. In gold, colour-coded t-values for each contrast thresholded at a significance level of p < 0.001, uncorrected and without masking. Results are overlaid on a Montreal Neurological Institute (MNI) brain. Note that although the significant clusters in (a), (c) and (e) all fall within the Automated Anatomical Labeling (AAL) definition of precuneus, the peak coordinates are different (see Table S10, Bi, Di, Ei). **(b, d, f)** Mean functional activity extracted from clusters significant at p_FWE_ < 0.05 shown in (a, c, e). The scatterplots are presented for visualisation purpose only and should not be used for statistical inference. (b) Red dots represent group mean ±SEM. Grey lines represent individual subjects. (d, f) Each data point represents a single participant, grey bands represent 95% confidence intervals. Pearson correlation effect sizes are shown. arb. u.: arbitrary units; S2-4: Session 2-4; n = 28 for (A-D), n = 21 for (E-F).

Next, we looked for a relationship between post-sleep performance improvements and brain activity changes, as shown previously^13,23,24^. Thus, we analysed the same [cued > uncued] contrast again, both for each session and for changes between sessions (S1 < S2, S2 < S3, S1 < S3). This time we included three regressors in separate models: behavioural cueing benefit at S2, S3 and S4 for the BH dataset. All clusters significant at p_FWE_ < 0.05 are reported in Table S10 D-F. As expected, when examining the cueing benefit at S2 as a regressor [(cued > uncued) * cueing benefit at S2] we found a TMR-related functional increase at S2 in left dorsal-posterior precuneus (−4, −78, 46; peak T = 5.18, p = 0.009; Fig.5C-D, Table S10Di, Fig.S1B). Interestingly, a ventral subregion of left precuneus (−2, −54, 10) showed a decreased response to TMR from S2 to S3 when considering behavioural cueing benefit at S4 as a regressor in the [Δ(cued > uncued) from S2 to S3] contrast (peak T = 6.66, p = 0.002; Fig.5E-F, Table S10Ei, Fig.S1C). Taken together, these results suggest that activity in dorsal precuneus 24 h post-encoding predicts behavioural effects of cueing in the short-term. However, longer-term benefits of cueing appear to be associated with an eventual decrease in response of the ventral precuneus.

In addition to precuneus, the cueing benefit at S4 regressor showed a positive relationship with TMR-related functional activity [(cued > uncued) * cueing benefit at S4] in the right postcentral gyrus at S3 (58, −18, 38; peak T = 5.50, p = 0.022; Fig.6A-B, Table S10Fi, Fig.S1D). This suggests that the way TMR impacts on activation of primary somatosensory cortex 10 days post-encoding may underpin the long-term behavioural effects of this cueing.

**Fig. 6.**
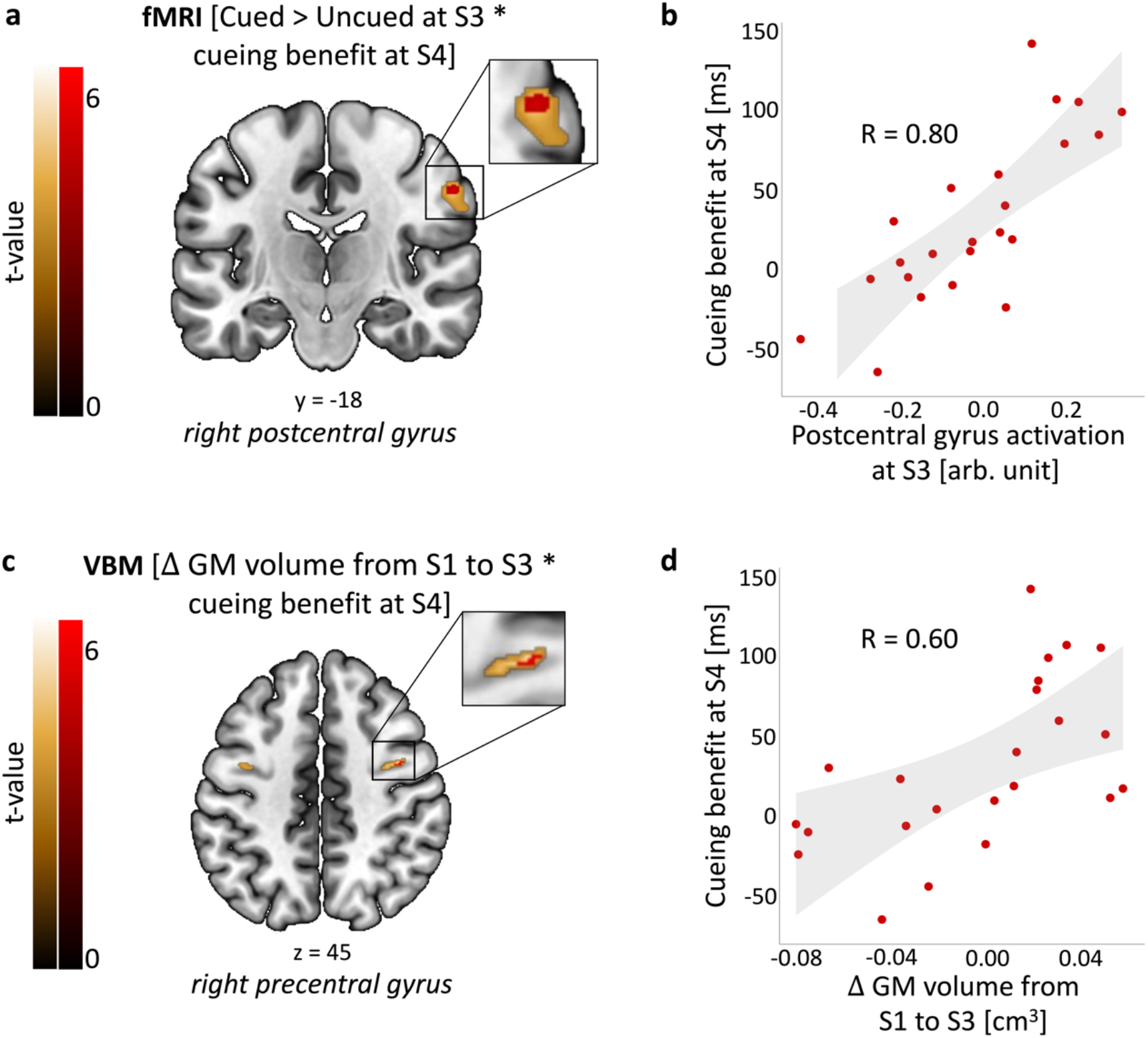
Functional activity and structural brain changes are associated with long-term cueing benefit. **(a-b)** Activity for the cued > uncued contrast in the right postcentral gyrus at S3 is positively associated with behavioural cueing benefit at S4. **(c-d)** Grey matter volume in the right precentral gyrus at S3 relative to S1 is positively associated with behavioural cueing benefit at S4. (**a, c**) Group level analysis. In red, colour-coded t-values for increased fMRI activity (a) and grey matter volume (c), both thresholded at a significance level of p_FWE_ < 0.05, corrected for multiple voxel-wise comparisons within a pre-defined ROI for bilateral sensorimotor cortex. In gold, colour-coded t-values for increased fMRI activity (a) and grey matter volume (c), both thresholded at a significance level of p < 0.001, uncorrected and without masking. Results are overlaid on a Montreal Neurological Institute (MNI) brain. Colour bars indicate t-values. **(b, d)** Mean functional activity (b) and grey matter volume (d) extracted from clusters significant at p_FWE_ < 0.05 shown in (a, c). The scatterplots are presented for visualisation purpose only and should not be used for statistical inference. Each data point represents a single participant, grey bands represent 95% confidence intervals. Pearson correlation effect sizes are shown. arb. u.: arbitrary units; GM: grey matter; S1-4: Session 1-4; n = 23.

### 2.5 TMR-RELATED STRUCTURAL PLASTICITY

To determine whether the behavioural effects of TMR were associated with volumetric changes, we performed voxel-based morphometry (VBM) analysis of the T1w scans. We thus carried out one-way t-tests on the subtraction images obtained for different time periods (S1 < S2, S2 < S3, S1 < S3), with behavioural regressors added as before. All clusters significant at p_FWE_ < 0.05 are reported in Table S11. When considering cueing benefit at S4 as a regressor we found a positive correlation with grey matter (GM) volume change from S1 to S3 [Δ GM volume from S1 to S3 * cueing benefit at S4] in the right precentral gyrus (42, −2, 45; peak T = 6.21, p = 0.020; Fig.6C-D, Table S11A, Fig.S2A). No correlations with volumetric changes were revealed in white matter. This is consistent with our fMRI finding (Fig.6A-B) and suggests that the time-dependent change in GM volume within a sensorimotor structure predicts long-term behavioural effects of cueing.

### 2.6 MEDIATION ANALYSIS

Having found that cueing benefit at S4 is associated with both TMR-related fMRI increase in right postcentral gyrus at S3 (Fig.6A-B) and GM volume increase from S1 to S3 in right precentral gyrus (Fig.6C-D), we set out to test for a mediation effect of the fMRI result on the relationship between the VBM result and behavioural cueing benefit. We reasoned that the early structural changes we observed may be shaping later functional changes in task-relevant areas, which in turn may be a direct driver of long-term cueing benefit. In line with our hypothesis, we found evidence for a significant mediation effect (Fig.7, 1-step mediation, β = 0.336, 95% CI = [0.146 – 0.604], indirect effect explaining 56% of total effect; Sobel test, Z = 2.32, p = 0.020). Furthermore, given that the direct effect was not significant (β = 0.597, p = 0.088; 95% CI: [-0.043 – 0.564]), we argue that the mediation is indirect-only^25^. Thus, our results suggest that GM change within the primary motor cortex drives the functional activity within the primary somatosensory cortex and thereby leads to the behavioural effects.

**Fig. 7.**
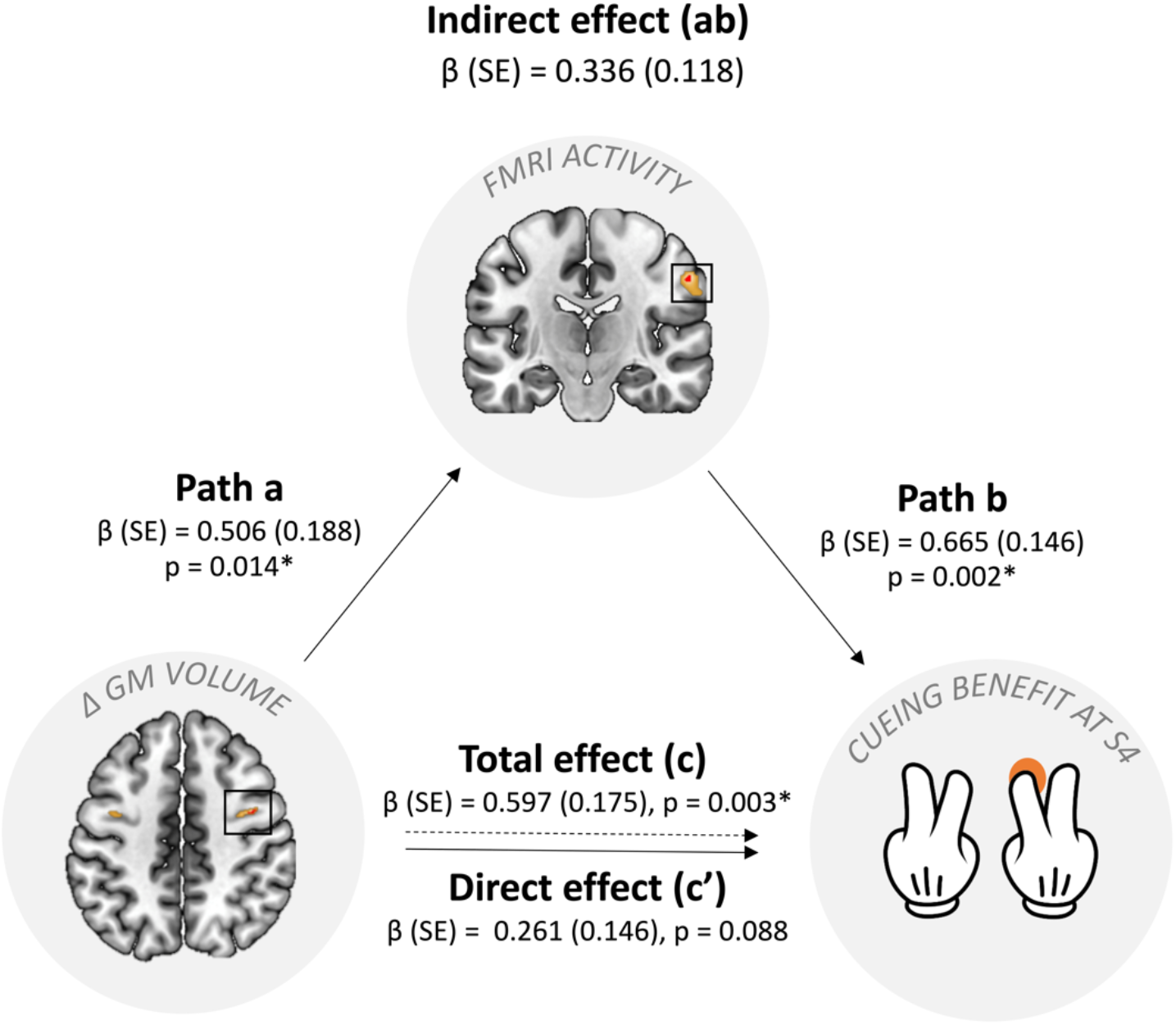
Results of mediation analysis. The diagram depicts standardized regression coefficients (β) for the relationship between the VBM result ([Δ GM volume from S1 to S3 * cueing benefit at S4] contrast) and cueing benefit at S4, mediated by the fMRI finding ([cued > uncued * cueing benefit at S4]). The associated standard error estimates (SE) are in parenthesis. Parameter estimates for the indirect effect were computed for 5,000 bootstrapped samples. Total effect explained 36% of the variance in behavioural cueing benefit and was significant (p = 0.003; 95% CI: [0.233-0.961]). Mediated model explained 68% of the variance. GM: grey matter; S4: Session 4. *p < 0.05. n = 23.

## 3 DISCUSSION

In this study we aimed to determine if repeated reactivation of a motor memory trace during sleep engenders learning-related changes within the PPC and sensorimotor areas. To this end, we tested the temporal dynamics of the TMR-related changes across structural, functional, electrophysiological, and behavioural measures. Firstly, we showed a main effect of TMR on the SRTT performance across all post-stimulation sessions, with the biggest difference between the cued and uncued sequence emerging 20 days poststimulation. In line with our hypothesis, dorsal precuneus showed a functional response that was both related to the manipulation and predicted its behavioural effects the next day. However, over time, both functional disengagement in the ventral subregion of precuneus and an increase in functional activity and volumetric grey matter in somatosensory and motor regions predicted the long-term behavioural benefit of our manipulation.

### TMR BENEFITS PROCEDURAL MEMORIES UP TO 20 DAYS POST-MANIPULATION

The strongest behavioural difference between the cued and uncued sequence occurred 20 days post-manipulation, suggesting that the benefits of cueing may last longer than previously believed. This is especially true given that neither object-location^13^ nor emotional^26^ memory seems to benefit from the manipulation even a week later. One-week-later effects of TMR have been reported for implicit biases^27^, but this failed to replicate^28^. Our prior work showed behavioural effects of TMR 10 days post-manipulation but not 6 weeks later^11^. Hence, the long-term effect of procedural memory cueing that we observe here appears to be the longest reported in the literature so far. Our findings also suggest that TMR starts a process which then unfolds over several weeks, gradually leading to the emergence of behavioural benefits over time.

### CUEING ENGAGES PRECUNEUS FUNCTIONALLY

Precuneus showed a TMR-dependent (cued > uncued) BOLD increase 24 h post-stimulation. Interestingly, this functional response predicted the behavioural cueing benefit at the same time point. These results combine to suggest that repeated reactivation of memory traces during sleep engages parts of the PPC in a behaviourally relevant manner. PPC has been identified as a hippocampus-independent memory store, whereby both hippocampal activity and hippocampal connectivity with PPC decreases after encoding, but (conversely) PPC activity increases over 24 h, as an independent memory representation builds up^29^. We believe that sleep plays a crucial role in this process and that the reactivation-mediated reorganisation of memories between the hippocampal-dependent short-term store and neocortex-dependent long-term store^4,30^ fosters engram development in the precuneus. Interestingly, prior observations of increased activity of cuneus, an area adjacent to precuneus, during *both* SRTT training *and* post-training sleep^31,32^ are consistent with our current findings. We speculate that memory reactivation could be taking place in cuneus, such that procedural memories are stored and processed in adjacent locations. Indeed, PPC has repeatedly been implicated in memory formation, retrieval, and storage^33–35^ and is traditionally associated with the motor system^36,37^. Our results suggest that precuneus may be particularly involved in early consolidation of memories that are reactivated during sleep.

Although we showed that TMR-related functional activity in precuneus is associated with behavioural cueing benefit 24 h post-manipulation, it is important to note that we find no group level evidence for behavioural cueing benefit at that time point. This could be due to the jittering of our TMR cues during sleep^11^. By randomising the inter-trial-interval between the TMR sounds we disrupted the temporal dynamics of sequence replay, decreasing the predictability of sequence elements. This may have delayed the impact of this manipulation on behaviour, such that behavioural impacts of TMR were not significant until 20 days post-manipulation. Even so, the absence of a TMR-related behavioural plasticity 10 days after cueing was unexpected given that cueing benefit was apparent at this time point in our prior study which used the same jittered TMR^11^. One possibility is that doing the task while lying down in the MRI scanner, and with somewhat clunky MR-safe button boxes, masked behavioural effects of TMR which would otherwise have been apparent. Indeed, the only session during which we observed a significant cueing benefit was the one which was performed online and in participants’ own homes using a computer keyboard.

Despite comparable performance on cued and uncued sequences 24 h post-TMR, participants exhibited distinct fMRI activation patterns for these two sequences at this time point. Indeed, we demonstrate that the dorsal subregion of precuneus is more functionally involved during production of the cued sequence 24 h post-TMR. Given that dorsal precuneus is specialised for somato-motor and visual-spatial processing^38^, this finding raises the possibility that visuomotor integration of the reactivated memories may underpin short-term cueing benefits, even if it is not enough to drive the behavioural plasticity. In turn, we show that the functional disengagement of ventral precuneus over time facilitates cueing benefits 20 days post-TMR. In contrast to the dorsal precuneus, its ventral subregion has been mostly implicated in episodic memory retrieval^38^. Hence, as ventral precuneus disengages, it perhaps allows other regions to take over, thus indirectly supporting longterm consolidation and behavioural performance.

### PLASTICITY WITHIN MOTOR REGIONS PREDICTS LONG-TERM CUEING BENEFITS

Our results suggest that a slowly evolving reorganisation of sensorimotor representations may underpin motor learning over a 20-day timescale. Specifically, we show that TMR-related functional activity in the right postcentral gyrus 10 days post-stimulation predicts behavioural benefits 20 days post-stimulation. Similarly, an increase in grey matter volume in the right precentral gyrus over the first 10 days post-stimulation predicts the behavioural benefits observed 20 days post-stimulation. In other words, both the functional activation and the volumetric grey matter increase in the sensorimotor cortex at 10 days post-TMR predict long-term cueing benefits.

Our mediation results further suggest that the functional activation of primary somatosensory cortex may be mediating a relationship between volumetric grey matter change in primary motor cortex and behavioural benefit from TMR. Indeed, these structures are known to be strongly linked^39^. In line with our findings, recent studies also demonstrate information flow from primary motor to primary sensory cortex, and vice versa (see ^40^ for review). Importantly, slowly evolving reorganisation of primary motor cortex after learning has been linked to long-term retention of motor skills^41–43^ and even referred to as the long-term motor engram^44,45^. Our findings suggest that memory reactivation during sleep may be crucial in the formation of this type of representation, reinforcing the long-suggested involvement of motor regions in sleep-dependent procedural memory consolidation^46,47^. Furthermore, we expand the existing literature by showing that task-related structural and functional plasticity can emerge weeks after a single night of cueing during sleep. Our data show that just one night of TMR can impact on brain plasticity in the long run, perhaps facilitating the development of a stable memory trace over subsequent nights of un-manipulated sleep.

Interestingly, both SRTT training and post-training REM sleep are linked to increased premotor activity, suggesting that REM sleep may be involved in the reactivation of motor memories^31^. Furthermore, an association has been observed between increased sensorimotor responses to the cued SRTT sequence and REM sleep^10^. Although the increased involvement of sensorimotor regions reported here was not related to the amount of time spent in REM sleep during the stimulation night, it was observed 10 days after encoding. In line with the suggestion that the delayed benefits of sleep may be due to processing in REM^48^ we propose that REM reactivation during post-TMR nights, rather than immediately after cueing, may contribute to the long-term effects of TMR. However, it is unknown whether sensorimotor reactivation is taking place in post-TMR REM and whether it contributes to the behavioural and neural benefits of the manipulation in the long run.

### THE ROLE OF N2 AND SLEEP SPINDLES

Finally, we recognise a fundamental similarity between the reactivation of memory traces via TMR and repeated encoding-retrieval episodes during wake, which have also been shown to engender rapid memory engram formation within precuneus^14^. Indeed, repeated retrieval is a powerful way to consolidate memories and shares a lot of parallels with offline reactivation^49^. However, in line with other studies^15^, we argue that the role of sleep goes beyond simply allowing opportunity for more rehearsal. Both N2^50,51^ and sleep spindles^20^ have been consistently implicated in motor sequence memory consolidation. Although we found no relationship between behavioural cueing benefit and either the time spent in N2 or spindle density, we did find a surge in spindle density during the cue period relative to the no-cue period. This is in line with our prior report^11^ and indicates that auditory cueing may elicit sleep spindles and thereby engender an immediate processing of memory traces^52,53^. Thus, our current results further support the relationship between spindles and procedural memory cueing.

### THE SEARCH FOR AN ENGRAM

A neuronal ensemble that holds a representation of a stable memory is known as an engram^54^. The term ‘engram’ also refers to the physical brain changes that are induced by learning and that enable memory recall^55^. Due to their widely distributed and dynamic nature, engrams have long remained elusive. However, recent technological advances allow us to study memory engrams in humans^55,56^. PPC, for instance, has received increasing attention in memory research^33^. Precuneus is a subregion of PPC and has been shown to undergo learning-dependent plasticity, fulfilling all criteria for a memory engram^14^. These defining criteria require an engram to (1) emerge as a result of encoding and reflect the content of the encoded information, (2) engender a persistent, physical change in the underlying substrate that (3) enables memory retrieval, and (4) exists in a dormant or inactive state, i.e., between encoding and retrieval processes^55^. Evidence for a relationship between engram formation and memory reactivation during sleep has so far been lacking. While previous literature suggests that changes in the precuneus alone fulfil all proposed criteria for an engram^14^, our data show no TMR-related structural changes in this region, and thus fail to fulfil criterion 2. This could be due to our use of different MRI modalities (i.e., structural rather than microstructural MRI as in ^14^). Nevertheless, if our results are considered collectively across regions, we can argue that they do fulfil the criteria for an engram. Specifically, we find that memory representations stored in precuneus and postcentral gyrus reflect the encoded information (criterion 1), and enable memory recall (criterion 3), whereas the precentral gyrus undergoes structural changes (criterion 2) over a relatively long period of time (criterion 4). Thus, our results suggest that memory reactivation during sleep could support the development and evolution of an engram that encompasses several cortical areas.

## 4 CONCLUSION

We show that the behavioural benefits of procedural memory cueing in NREM sleep develop over time and emerge 20 days post-encoding. Increased TMR-related engagement of dorsal precuneus underpins the shortterm effects of stimulation (over 24 hours), whereas sensorimotor regions support the long-term effects (over 20 days). These results advance our understanding of the neural changes underpinning long-term offline skill consolidation. They also shed new light on the TMR-induced processes that unfold over several nights after auditory cueing.

## 5 MATERIALS AND METHODS

### 5.1 PARTICIPANTS

A pre-study questionnaire was used to exclude subjects with a history of drug/alcohol abuse, psychological, neurological or sleep disorders, hearing impairments, recent stressful life event(s) or regular use of any medication or substance affecting sleep. Participants were required to be right-handed, non-smokers, have regular sleep pattern, normal or corrected-to-normal vision, no prior knowledge of the tasks used in the study, and no more than three years of musical training in the past five years as musical training has previously been shown to affect procedural learning^57,58^. None of the participants reported napping regularly, working night shifts or travelling across more than two time-zones one month prior to the experiment. 33 volunteers fulfilled all inclusion criteria and provided an informed consent to participate in the study, which was approved by the Ethics Committee of the School of Psychology at Cardiff University (ethics number EC.19.06.11.5651R3A2) and performed in accordance with the Declaration of Helsinki. All participants agreed to abstain from extreme physical exercise, napping, alcohol, caffeine, and other psychologically active food from 24 h prior to each experimental session. Finally, before their first session, participants were screened by a qualified radiographer from Cardiff University to assess their suitability for MRI and signed an MRI screening form prior to each scan.

Three participants had to be excluded from all analyses due to: technical issues (n = 1), voluntary withdrawal (n = 1), and low score on the handedness questionnaire (indicating mixed use of both hands), combined with a positive slope of learning curve during the first session (indicating lack of sequence learning before sleep) (n = 1). Hence, the final dataset included 30 participants (16 females, age range: 18 – 23 years, mean ±SD: 20.38 ± 1.41; 14 males, age range: 19 – 23 years, mean ±SD: 20.43 ± 1.16). However, due to the COVID-19 outbreak, six participants were unable to complete the study, missing all data from either one (n = 1) or two (n = 5) sessions. Hence, n = 25 for all data collected during S3 and n = 24 for S4. The final dataset included one participant who could not physically attend S3. They performed the SRTT online, but their MRI data (functional, fMRI and structural, T1w) could not be collected and therefore the sample size for the MRI analyses of S3 had to be further decreased by one. Two additional participants were excluded from the fMRI analysis of S2 due to MRI gradient coil damage during fMRI acquisition (n = 1) and failure to save the fMRI data (n = 1). Hence, the final sample size for fMRI was n = 30 for S1, n = 28 for S2 and n = 24 for S3, whereas the final sample size for analysis of T1w data was n = 30 for S1, n = 30 for S2 and n = 24 for S3. Finally, one participant had to be excluded from all the analyses concerning EEG due to substantial loss of data caused by failure of the wireless amplifier during the night. However, the TMR procedure itself was unaffected and therefore this participant was included in the behavioural and MRI analyses. A flowchart of participants included and excluded from the different analyses is presented in Fig.S3.

### 5.2 STUDY DESIGN

The experiment consisted of four sessions (Fig.1A), all scheduled for ~8 pm. Upon arrival for the first session, participants completed Pittsburgh Sleep Quality Index (PSQI)^59^ to examine their sleep quality over the past month and Stanford Sleepiness Questionnaire (SQ)^60^ to assess their current level of alertness. A short version of the Edinburgh Handedness Inventory (EHI)^61^ was also administered to confirm that all subjects were righthanded before the learning session took place. Due to time constraints at the MRI scanner the learning session had to be split into two parts. The first half of the SRTT blocks (24 sequence blocks) were performed in a 0T Siemens ‘mock’ scanner which also helped to acclimate subjects to the scanner environment. The second half of the SRTT blocks (24 sequence blocks + 4 random blocks) was performed in a 3T Siemens MRI scanner during fMRI acquisition and used for functional data analysis. fMRI acquisition was preceded by a structural scan (T1w) and followed by a B0 fieldmap (see section *2.6* *MRI data acquisition*). Once outside the MRI scanner, participants were asked to prepare themselves for bed. They were fitted with an EEG cap and were ready for bed at ~11 pm. During N2 and N3 sleep stages, tones associated with one of the SRTT sequences were replayed to the participants via speakers (Harman/Kardon HK206, Harman/Kardon, Woodbury, NY, USA) to trigger reactivation of the SRTT memories associated with them. Participants were woken up after, on average, 8.81 ±0.82 h in bed and had the EEG cap removed before leaving the lab.

We asked participants to come back for the follow-up sessions 23-26 h (session 2, S2), 10-14 days (session 3, S3) and 16-21 days (session 4, S4) after S1. During S2, participants were asked to indicate if they remember hearing any sounds during the night in the lab. S2 and S3 lasted ~2 h each and both involved the SQ and an MRI scan, during which a structural scan was acquired. This was followed by the SRTT re-test, with the first half of the SRTT blocks (24 sequence blocks + 4 random blocks) performed during the fMRI acquisition and the second half (24 sequence blocks + 4 random blocks) in the mock scanner. Note that the order of scanners (3T vs 0T) was flipped from S1 to S2 and S3 for the functional and structural assessment to occur as close to the TMR session as possible. S4 took place either in the lab or online, depending on the severity of COVID-19 restrictions at the time. During S4, SQ was delivered as before, together with the SRTT (one run, 48 sequence blocks + 4 random blocks) and an explicit memory task. Upon completion of each session, participants were informed about the upcoming SRTT re-tests as this has been shown to enhance post-learning sleep benefits^62^.

For offline data collection, the SRTT (S1-S3) was back projected onto a projection screen situated at the end of the MRI/mock scanner and reflected into the participant’s eyes via a mirror mounted on the head coil; the questionnaires and the SRTT (S4) were presented on a computer screen with resolution 1920 x 1080 pixels, and the explicit memory task was completed with pen and paper. SRTT was presented using MATLAB 2016b (The MathWorks Inc., Natick, MA, USA) and Cogent 2000 (developed by the Cogent 2000 team at the Functional Imaging Laboratory and the Institute for Cognitive Neuroscience, University College, London, UK; http://www.vislab.ucl.ac.uk/cogent.php); questionnaires were presented using MATLAB 2016b and Psychophysics Toolbox Version 3^63^.

For online data collection, SRTT (S4) was coded in Python using PsychoPy 3.2.2.^64^ and administered through the Pavlovia online platform (https://pavlovia.org/); questionnaires were distributed via Qualtrics software^65^, and the explicit memory task was sent to the participants as a .pdf document which they were asked to edit according to the instructions provided.

### 5.3 EXPERIMENTAL TASKS

#### 5.3.1 MOTOR SEQUENCE LEARNING – THE SERIAL REACTION TIME TASK (SRTT)

The SRTT (Fig.1B) was used to induce and measure motor sequence learning. It was adapted from^66^, as described previously^11^. SRTT consists of two 12-item sequences of auditorily and visually cued key presses, learned by the participants in blocks. The task was to respond to the stimuli as quickly and accurately as possible, using index and middle fingers of both hands. The two sequences – A (1–2–1–42–3–4–1–3–2–4–3) and B (2–4–3–2–3–1–4–2–3–1–4–1) – were matched for learning difficulty, they did not share strings of more than four items and contained items that were equally represented (three repetitions of each). Each sequence was paired with a set of 200 ms-long tones, either high (5^th^ octave, A/B/C#/D) or low (4^th^ octave, C/D/E/F) pitched, that were counterbalanced across sequences and participants. For each item/trial, the tone was played with simultaneous presentation of a visual cue in one of the four corners of the screen. Visual cues consisted of neutral faces and objects appearing in the same location regardless of the sequences (1 – top left corner = male face, 2 – bottom left corner = lamp, 3 – top right corner = female face, 4 – bottom right corner = water tap). Participants were told that the nature of the stimuli (faces/objects) was not relevant for the study. Their task was to press the key on the keyboard (while in the sleep lab or at home) or on an MRI-compatible button pad (2-Hand system, NatA technologies, Coquitlam, Canada) (while in the MRI/mock scanner) that corresponded to the position of the picture as quickly and accurately as possible: 1 = left shift/left middle finger button; 2 = left Ctrl/left index finger button; 3 = up arrow/right middle finger button; 4 = down arrow/right index finger button. Participants were instructed to use both hands and always keep the same fingers on the appropriate response keys. The visual cue disappeared from the screen only after the correct key was pressed, followed by a 300 ms interval before the next trial.

There were 24 blocks of each sequence (a total of 48 sequence blocks per session). The block type was indicated with ‘A’ or ‘B’ displayed in the centre of the screen. Each block contained three sequence repetitions (36 items) and was followed by a 15 s pause/break, with reaction time and error rate feedback. Blocks were interleaved pseudo-randomly with no more than two blocks of the same sequence in a row. Participants were aware that there were two sequences but were not asked to learn them explicitly. Block order and sequence replayed were counterbalanced across participants.

During each run of the SRTT, sequence blocks A and B were followed by 4 random blocks except for the first half of S1 (to avoid interrupted learning). Random blocks were indicated with ‘R’ appearing in the centre of the screen and contained pseudo-randomised sequences. For these, visual stimuli were the same and tones matched sequence A tones for half of them (Rand_A) and sequence B tones for the other half (Rand_B). Blocks Rand_A and Rand_B were alternated, and each contained random sequences constrained by the following criteria: (1) cues within a string of 12 items were equally represented, (2) the same cue did not occur in consecutive trials, (3) the sequence did not share more than four cues in a row with either sequence A or B.

#### 5.3.2 EXPLICIT MEMORY TASK

Explicit memory of the SRTT was assessed by a free recall test administered at the end of the study (S4). Participants were provided with printed screenshots of sequence A and sequence B trials, but the visual cues were removed. They were instructed to mark the order of each sequence by drawing an ‘X’ to indicate cue location.

### 5.4 EEG DATA ACQUISITION

EEG data was acquired with actiCap slim active electrodes (Brain Products GmbH, Gilching, Germany). 62 scalp electrodes were embedded within an elastic cap (Easycap GmbH, Herrsching, Germany), with the reference electrode positioned at CPz and ground at Afz. Electromyogram (EMG) signals were recorded from two electrodes placed on the chin, whereas the electrooculogram (EOG) was collected from two electrodes placed below the left eye and above the right eye. EEG electrodes layout is presented in Fig.S4. Elefix EEG-electrode paste (Nihon Kohden, Tokyo, Japan) was applied on each electrode for stable attachment and Super-Visc high viscosity electrolyte gel (Easycap GmbH) was used to keep impedance below 25 kOhm. Signals were amplified with either two BrainAmp MR plus EEG amplifiers or LiveAmp wireless amplifiers and recorded using BrainVision Recorder software (all from Brain Products GmbH).

### 5.5 TMR DURING NREM SLEEP

The TMR protocol was administered as before^11^, using MATLAB 2016b and Cogent 2000. Briefly, tones associated with either sequence A or B (counterbalanced across participants) were replayed to the participants during stable N2 and N3 (Fig.1C). Replay blocks contained one repetition of a sequence (i.e., 12 sounds) and were followed by 20 s of silence. The inter-trial interval between individual sounds was jittered between 2,500 and 3,500 ms. Volume was adjusted for each participant to prevent arousal. However, upon leaving the relevant sleep stage, replay was paused and resumed only when stable N2 or N3 was observed. TMR was performed until ~1,000 trials were delivered in N3. On average, 1385.20 ±305.53 sounds were played.

### 5.6 MRI DATA ACQUISITION

Magnetic resonance imaging (MRI) was performed at Cardiff University Brain Imaging Centre (CUBRIC) with a 3T Siemens Connectom scanner (maximum gradient strength 300 mT/m). All scans were acquired with a 32-channel head-coil and lasted ~1 h in total each, with whole-brain coverage including cerebellum. Apart from the T1w and fMRI scans, the MRI protocol also included multi-shell Diffusion-Weighted Imaging (DWI) and mcDESPOT acquisitions, but these are not discussed here.

#### 5.6.1 T1-WEIGHTED IMAGING

A high resolution T1w anatomical scan was acquired with a 3D magnetization-prepared rapid gradient echoes (MPRAGE) sequence (2300 ms repetition time [TR]; 2 ms echo time [TE]; 857 ms inversion time [TI]; 9° flip angle [FA]; bandwidth 230 Hz/Pixel; 256 mm field-of-view [FOV]; 256 x 256 voxel matrix size; 1 mm isotropic voxel size; 1 mm slice thickness; 192 sagittal slices; parallel acquisition technique [PAT] with in-plane acceleration factor 2 (GRAPPA); anterior-to-posterior phase-encoding direction; 5 min total acquisition time [AT]) at the beginning of each scanning session.

#### 5.6.2 FUNCTIONAL MRI

Functional data were acquired with a T2*-weighted multi-band echo-planar imaging (EPI) sequence (2,000 ms TR; 35 ms TE; 75° FA; bandwidth 1976 Hz/Pixel; 220 mm FOV; 220 x 220 voxel matrix size; 2 mm isotropic voxel size; 2 mm slice thickness; 87 slices with a ~25° axial-to-coronal tilt from the anterior – posterior commissure (AC-PC) line and interleaved slice acquisition; PAT 2 (GRAPPA); multi-band acceleration factor [MB] 3; anterior-to-posterior phase-encoding direction; maximum 24 min AT and 720 scans; because the task was self-paced the exact AT and the number of scans differed between participants). Each fMRI acquisition was preceded by dummy scans to allow for saturation of the MR signal before the start of the task. Due to the nature of the task, the fMRI paradigm followed a block design consisting of sequence and random blocks (self-paced), alternating with rest blocks (15 s) (see *section 2.3.1 Motor sequence learning* – *the serial reaction time task* (*SRTT*)). Presentation of the first stimulus in a block was synchronised with the scanner’s trigger signal sent upon acquisition of every fMRI volume. Thus, the beginning of the task (i.e., the first stimulus of the first block) was triggered by the first fMRI volume acquisition and for that reason the initial volumes did not have to be discarded. No online motion correction was applied.

#### 5.6.3 B0 FIELDMAP

B0-fieldmap was acquired to correct for distortions in the fMRI data caused by magnetic field (i.e., B0) inhomogeneities (465 ms TR; 4.92 ms TE; 60° FA; bandwidth 290 Hz/Pixel; 192 mm FOV; 192 x 192 voxel matrix size; 3 mm isotropic voxel size; 3 mm slice thickness; 44 slices with a ~25° axial-to-coronal tilt from the AC-PC line and interleaved slice acquisition; 1 average; anterior-to-posterior phase-encoding direction; 1 min AT).

### 5.7 DATA ANALYSIS

#### 5.7.1 BEHAVIOURAL DATA

##### 5.7.1.1 SRTT: REACTION TIME

SRTT performance was measured using mean reaction time per block of each sequence (cued and uncued). Both hands (BH) dataset contained all SRTT trials within each block, except for those with reaction time exceeding 1,000 ms. Trials with incorrect button presses prior to the correct ones were included in the analysis. BH dataset was divided into the right hand (RH) dataset and left hand (LH) dataset, where each contained only the trials performed with the dominant or non-dominant hand, respectively. For each sequence within a given dataset, the mean performance on the 4 target blocks (brown and grey vertical bars in Fig.2A) was subtracted from the mean performance on the 2 random blocks. This allowed us to separate sequence learning from sensorimotor mapping and thus obtain a measure of ‘sequencespecific skill’ (SSS). The target blocks were the first 4 sequence blocks, used to calculate early SSS, and the last 4 sequence blocks, used to calculate late SSS, as illustrated below:

1. Early SSS = mean (random blocks) – mean (first 4 sequence blocks)
2. Late SSS = mean (random blocks) – mean (last 4 sequence blocks)

Finally, to obtain a single measure reflecting the effect of TMR on the SRTT performance we calculated the difference between the SSS of the cued and uncued sequence and refer to it as the ‘cueing benefit’.

##### 5.7.1.2 QUESTIONNAIRES

PSQI global scores were determined in accordance with the original scoring system^59^ (Buysse et al., 1989). Answers to the short version of the EHI were scored as in ^61^ and used to obtain laterality quotient for handedness. For results, see *Supplementary Notes: Questionnaires*.

##### 5.7.1.3 EXPLICIT MEMORY

Responses on the explicit memory task were considered correct only if they were in the correct position within the sequence and next to at least one other correct item, hence reducing the probability of guessing^66^. The number of items guessed by chance was determined for each participant by taking an average score of 10 randomly generated sequences. To test if the explicit memory was formed, the average chance level across all participants was compared with the average number of correct items for each sequence. For results, see *Supplementary Notes: Explicit memory task* and Fig.S5.

#### 5.7.2 EEG DATA ANALYSIS

All EEG data were analysed in MATLAB 2018b using FieldTrip Toolbox^67^.

##### 5.7.2.1 SLEEP SCORING

EEG signal recorded throughout the night at eight scalp electrodes (F3, F4, C3, C4, P3, P4, O1, O2), two EOG and two EMG channels was pre-processed and re-referenced from CPz to the mastoids (TP9, TP10). For two participants, the right mastoid channel (TP10) was deemed noisy through visual inspection and had to be interpolated based on its triangulation-based neighbours (TP8, T8, P8), before it could be used as a new reference. The data was scored according to the AASM criteria^68^ by two independent sleep scorers who were blind to the cue presentation periods. Sleep scoring was performed using a custom-made interface (https://github.com/mnavarretem/psgScore).

##### 5.7.2.2 SPINDLES ANALYSIS

The relationship between sleep spindles and behavioural measures was assessed using 8 electrodes located over motor areas: FC3, C5, C3, C1, CP3, FC4, C6, C4, C2, CP4. However, for visualisation purposes (Fig.4A), the remaining electrodes in the International 10-20 EEG system were also analysed as described below. First, raw data from these channels were down-sampled to 250 Hz (for them to be comparable between the two EEG data acquisition systems) and filtered by Chebyshev Type II infinite impulse response (IIR) filter (passband: f = [0.3 – 35] Hz; stopband: f < 0.1 Hz & f > 45 Hz). All channels were visually inspected, and the noisy ones were interpolated via triangulation of their nearest neighbours. As a final pre-processing step, we re-referenced the data from CPz to the mastoids (TP9, TP10). A spindle-detection algorithm^69^ was then employed to automatically identify sleep spindles (11 – 16 Hz). Briefly, the data were filtered in a sigma band by the IIR filter (passband: f = [11 – 16] Hz; stopband: f < 9 Hz & f > 18 Hz) and the root mean squared (RMS) of the signal was computed using a 300 ms time window. Any event that surpassed the 86.64 percentile (1.5 SD, Gaussian distribution) of the RMS signal was considered a candidate spindle. To fit the spindle detection criteria^70^, only the events with unimodal maximum in the 11 – 16 Hz frequency range in the power spectrum, duration between 0.5 and 2.0 s and at least 5 oscillations were regarded as sleep spindles^69^.

Any identified spindles that fell (partly or wholly) within a period that had been previously marked as an arousal during sleep scoring were removed. The remaining spindles were separated into those that fell within the cue and no-cue periods. We define the cue period as the 3.5 s time interval after the onset of each tone. Since 3.5 s was the longest inter-trial interval allowed, the cue period essentially covered the time interval from the onset of the first tone in a sequence to 3.5 s after the onset of the last one. In turn, the no-cue period covered the time interval between sequences, i.e., from 3.5 to 20.0 s after the onset of the last tone in a sequence. If a spindle fell between the cue and no-cue period, that spindle was removed from further analysis. Thus, only spindles that fell wholly within the cue or no-cue period were included in the analysis.

Spindle density was calculated by dividing the number of spindles at each electrode and in each period of interest (cue period during target sleep stage, no-cue period during target sleep stage) by the duration (in minutes) of that period.

#### 5.7.3 MRI DATA ANALYSIS

MRI data were pre-processed using Statistical Parametric Mapping 12 (SPM12; Wellcome Trust Centre for Neuroimaging, London, UK), running under MATLAB 2018b.

##### 5.7.3.1 FMRI

###### PRE-PROCESSING

Functional data pre-processing consisted of (1) B0-fieldmap correction using SPM’s fieldmap toolbox^71^; (2) realignment to the mean of the images using a least-squares approach and 6 parameter rigid body spatial transformation to correct for movement artifact^72^; (3) co-registration with the participants’ individual structural image using rigid body model^73^; (4) spatial normalisation to Montreal Neurological Institute brain (MNI space) via the segmentation routine and resampling to 2 mm voxels with a 4^th^ degree B-spline interpolation^74^; (5) smoothing with 8 mm full-width half maximum (FWHM) Gaussian kernel in line with the literature^10^. All steps were performed as implemented in SPM12. B0-fieldmap correction step was omitted for one participant (n = 1) due to technical issues during B0-fieldmap acquisition. No scans had to be excluded due to excessive movement (average translations < 3.3 mm, average rotations < 0.03°).

###### SINGLE SUBJECT LEVEL ANALYSIS

Subject-level analysis of the fMRI data was performed using a general linear model (GLM)^75^, constructed separately for each participant and session. Each block type (cued sequence, uncued sequence, cued random, uncued random) as well as the breaks between the blocks were modelled as five separate, boxcar regressors; button presses were modelled as single events with zero duration. All of these were temporally convolved with a canonical hemodynamic response function (HRF) model embedded in SPM, with no derivatives. To control for movement artifacts, the design matrix also included six head motion parameters, generated during realignment, as non-convolved nuisance regressors. A high-pass filter with a cut-off period of 128 s was implemented in the matrix design to remove low-frequency signal drifts. Finally, serial correlations in the fMRI signal were corrected for using a first-order autoregressive model during restricted maximum likelihood (REML) parameter estimation. Contrast images were obtained for each block type of interest ([cued sequence] and [uncued sequence]), as well as for the difference between the two ([cued > uncued]). The resulting parameter images, generated per participant and per session using a fixed-effects model, were then used as an input for the group-level (i.e., random effects) analysis. Contrast images for the difference between sequence and random blocks were not generated due to the unequal number of each block type performed in the scanner (2 random blocks vs 24 sequence blocks, per session). This, however, was in accordance with the literature^10^.

##### 5.7.3.2 VBM

###### PRE-PROCESSING

Pre-processing of T1w images was performed in keeping with ^76^ recommendations. Images were first segmented into three tissue probability maps (grey matter, GM; white matter, WM; cerebrospinal fluid, CSF), with two Gaussians used to model each tissue class, very light bias regularisation (0.0001), 60 mm bias FWHM cut-off and default warping parameters^74^. Spatial normalisation was performed with DARTEL^77^, where the GM and WM segments were used to create customized tissue-class templates and to calculate flow fields. These were subsequently applied to the native GM and WM images of each subject to generate spatially normalised and Jacobian scaled (i.e., modulated) images in the MNI space, resampled at 1.5 mm isotropic voxels. The modulated images were smoothed with an 8 mm FWHM Gaussian kernel, in line with the fMRI analysis. To account for any confounding effects of brain size we estimated the total intracranial volume (ICV) for each participant at each time point by summing up the volumes of the GM, WM, and CSF probability maps, obtained through segmentation of the original images. The GM and WM images were then proportionally scaled to the ICV values by means of dividing intensities in each image by the image’s global (i.e., ICV) value before statistical comparisons.

#### 5.7.4 STATISTICAL ANALYSIS

All tests conducted were two-tailed, with the significance threshold set at 0.05. For behavioural and EEG data analyses, normality assumption was checked using Shapiro-Wilk test. To compare two related samples, we used paired-samples t-test or Wilcoxon signed-rank test, depending on the Shapiro-Wilk test result. Results are presented as mean ± standard error of the mean (SEM), unless otherwise stated.

##### 5.7.4.1 BEHAVIOURAL DATA

Statistical analysis of the behavioural data was performed in R^78^ or SPSS Statistics 25 (IBM Corp., Armonk, NY, USA) as before^11^. Each dataset (LH, RH, BH) was analysed separately.

To assess the relationship between TMR, SSS and Session we used linear mixed effects analysis performed on S2-S4, using lme4 package^79^ in R. We chose linear mixed effects analysis to avoid listwise deletion due to missing data at S3 and S4 and to account for the non-independence of multiple responses collected over time. TMR and Session were entered into the model as categorical (factor) fixed effects without interaction and random intercept was specified for each subject. The final models fitted to the BH, LH and RH datasets were as follows:

*> model = lmer(early SSS ~ Session + TMR + (1|Participant), data=dataset)*
*> model = lmer(late SSS ~Session + TMR + (1|Participant), data=dataset)*

To test for the effect of hand, LH and RH datasets were combined and ‘hand’ (factor) was added as an additional fixed effect:

*> model = lmer(early SSS ~ Session + TMR + Hand + (1|Participant), data=dataset)*
*> model = lmer(late SSS ~ Session + TMR + Hand + (1|Participant), data=dataset)*

Finally, to explore how the TMR effect evolves from S2 to S4, we entered cueing benefit (calculated using the late SSS data given no TMR effect on the early SSS) as the dependent variable and the number of days post-TMR (‘time’, integer) as a fixed effect in the following model:

*> model = lmer(CueingBenefit ~ Time + (1|Participant), data=dataset)*

To test for the effect of hand, LH and RH datasets were combined as before:

*> model = lmer(CueingBenefit ~ Time + Hand + (1|Participant), data=dataset)*

Likelihood ratio tests comparing the full model against the model without the effect of interest were performed using the ANOVA function in R to obtain p-values. Post-hoc pairwise comparisons were conducted using the *emmeans* package^80^ in R and corrected for multiple comparisons with Holm’s method. Effect sizes were calculated with the *emmeans* package as well.

##### 5.7.4.2 EEG DATA

Statistical analysis of the EEG data was performed in R^78^ or SPSS Statistics 25 (IBM Corp., Armonk, NY, USA). Each stimulation period (cue vs no-cue) and sleep stage (N2, N3, N2 and N3 combined) was analysed separately.

Correlations between our behavioural measures and EEG results were assessed with Pearson’s correlation or Spearman’s Rho (depending on the Shapiro-Wilk test result), using *cor.test* function in the R environment. Any datapoint that was both (1) more than 1.5 IQRs below the first quartile or 1.5 IQRs above the third quartile, and (2) deemed an outlier through visual inspection, was removed from the dataset prior to correlational analysis. False discovery rate (FDR) correction was used to correct for multiple correlations (q < 0.05)^81^. FDR corrections were based on 3 correlations, given the 3 experimental sessions of interest (S2, S3, S4).

##### 5.7.4.3 MRI DATA

Group level analysis of the MRI data was performed either in a Multivariate and Repeated Measures (MRM) toolbox (https://github.com/martynmcfarquhar/MRM) or in SPM12, both running under MATLAB 2018b. All tests conducted were two-tailed, testing for both positive and negative effects. Results were voxel-level corrected for multiple comparisons by family wise error (FWE) correction for the whole brain and for the pre-defined anatomical regions of interest (ROI), with the significance threshold set at p_FWE_ < 0.05. For the analysis performed in MRM, p-values were derived from 1,000 permutations, with Wilk’s lambda specified as the test statistic. Pre-defined ROI included (1) bilateral precuneus, (2) bilateral hippocampus and parahippocampus, (3) bilateral dorsal striatum (putamen and caudate), (4) bilateral cerebellum, (5) bilateral sensorimotor cortex (precentral and postcentral gyri). All ROI were selected based on their known involvement in sleep-dependent procedural memory consolidation^23,24,46,47^ and memory reactivation^6,10,12,14,31^. A mask for each ROI was created using an Automated Anatomical Labeling (AAL) atlas in the Wake Forest University (WFU) PickAtlas toolbox^82^. Anatomical localisation of the significant clusters was determined with the automatic labelling of MRIcroGL (https://www.nitrc.org/projects/mricrogl/) based on the AAL atlas. All significant clusters are reported in the tables, but only those with an extent equal to or above 5 voxels are discussed in text and presented in figures.

###### FMRI DATA

To test the effect of TMR on the post-stimulation sessions (S2-S3), one-dimensional contrast images for the [cued] and [uncued] blocks of each session were entered into a repeated-measures TMR-by-Session ANOVA performed in the MRM toolbox.

To compare functional brain activity during the cued and uncued sequence we carried out one-way t-tests on the [cued > uncued] contrast for S2 (n = 28) and S3 (n = 24) in SPM12. To determine the relationship between the TMR-related functional activity and other factors, we included the behavioural cueing benefit for the BH dataset at different time points (S2, S3, S4) as covariates in separate comparisons. Finally, to investigate fMRI changes over time, images from consecutive sessions were subtracted from one another, resulting in three subtraction images per subject (S1-S2, n = 28; S2-S3, n = 22; S1-S3, n = 24). We then performed the one-way t-tests as before (either with or without a covariate of interest).

###### VBM DATA

Group-level analysis of the structural images was performed separately for GM and WM. First, the preprocessed and proportionally scaled images from consecutive sessions were subtracted from one another as for fMRI (S1-S2, n = 30; S2-S3, n = 24; S1-S3, n = 24). To determine the relationship between the longitudinal brain changes and other factors, one-sample t-tests were computed in SPM12, with covariates of interest added one at a time. The covariates of interest were the behavioural cueing benefit for the BH dataset at a chosen time point (S2, S3, S4). Sex was always specified as a covariate of no interest (nuisance covariate) to control for differences between males and females. Finally, the SPM12 tissue probability maps of GM and WM were thresholded at 50% probability and the resulting binary masks were used in the analyses of the relevant tissue^83^.

##### 5.7.4.4 MEDIATION ANALYSIS

To determine the relationship between VBM, fMRI and behavioural measures, we conducted mediation effect parametric mapping based on a standard 3-variable path model^84^ with a bootstrap test (5,000 bootstrapped samples) for the statistical significance of the indirect effect^85^. The mediation analysis was performed using *PROCESS* macro for SPSS^86^. Cueing benefit at S4 was entered as the dependent variable, the average GM volume within the significant cluster from the [[Δ GM volume from S1 to S3 * cueing benefit at S4] contrast (i.e., precentral gyrus, p_FWE_ < 0.05, ROI corrected) was entered as the independent variable and the functional activity within the significant cluster from the [(cued > uncued) * cueing benefit at S4] contrast (i.e., postcentral gyrus, p_FWE_ < 0.05, ROI corrected) was entered as the mediator. All variables were standardised prior to entering in the mediation analysis. The percentage mediation denotes the ratio between the indirect effect and the total effect. Finally, a Sobel test^87^ was conducted to further demonstrate the significance using an interactive calculator tool for mediation test^88,89^.

#### 5.7.5 RESULTS PRESENTATION

Plots displaying behavioural results, pairwise comparisons and relationships between two variables were generated using *ggplot2* (version 3.3.0)^90^ in R. Fig.4A was generated using *ft_topoplotER* function in FieldTrip Toolbox^67^. Fig.1, Fig.7, Fig.S3 and Fig.S4 were created in Microsoft PowerPoint v16.53. MRI results are presented using MRIcroGL, displayed on the MNI152 standard brain (University of South Carolina, Columbia, SC), except Fig.S1 and Fig.S2 which were generated by SPM12 (Wellcome Trust Centre for Neuroimaging, London, UK).

## DATA AVAILABILITY

All data collected during the study, scripts that delivered experimental tasks and codes used to conduct the analyses are publicly available at: https://osf.io/y43sb/?view_only=8b18dd7984e94c629274bbf427fb90be.

## CODE AVAILABILITY

A custom-made interface used to perform sleep scoring can be accessed at: https://github.com/mnavarretem/psgScore.

## ACKNOWLEDGEMENTS

The authors would like to thank Holly Kings and Sofia Pereira for helpful comments and discussion on this manuscript, Chen Song and Marco Bigica for advice on the MRI analysis, as well as Chelsea Bryant and Joe Davis for support with participants’ recruitment and data collection. The authors are also grateful to Jennifer Roebber for sharing her SRTT script on Pavlovia and her help with Python coding. Finally, we thank all the participants for their time and commitment to the study. This work was supported by the ERC grant SolutionSleep, 681607, to PL.

## AUTHORS CONTRIBUTION STATEMENT

M.R. and P.A.L. conceived the study and designed the experiment, M.R., P.B. and M.E.A.A. carried it out; M.R., P.B and A.L performed the analysis; M.N. developed sleep scoring and EEG analysis algorithms; M.R wrote the manuscript with input from all co-authors; P.A.L. supervised the project and obtained funding.

