## Supplementary Materials for "Cueing motor memory reactivation during NREM sleep engenders learning-related changes in precuneus and sensorimotor structures"

### SUPPLEMENTARY NOTES: BASELINE SRTT PERFORMANCE

Before sleep, no difference was found between the average reaction time of the cued and uncued sequence for either both hands (BH, $t_{29} = -0.25$, $p = 0.801$), left hand (LH, $t_{29} = 0.27$, $p = 0.786$) or right hand (RH, $t_{29} = -0.50$, $p = 0.621$) (paired-samples t-tests) dataset. Similar results were obtained where comparing random sequences before sleep for all datasets (BH: $z = -0.57$, $p = 0.572$; LH: $z = -0.63$, $p = 0.530$; Wilcoxon signed-rank tests; RH: $t_{29} = -0.16$, $p = 0.872$; paired-samples t-test). Thus, any post-sleep difference between the sequences can be regarded as the effect of TMR. Furthermore, average reaction times before sleep were significantly shorter for the last 4 sequence blocks than for the random blocks, confirming that the participants learned both sequences during S1 (BH cued: $z = -4.74$, BH uncued: $z = -4.49$; LH cued: $z = -4.78$, LH uncued: $z = -4.33$; $p < 0.001$ for all comparisons, Wilcoxon signed-rank test; RH cued: $t_{29} = 7.21$, RH uncued: 6.49; $p < 0.001$ for all comparisons, paired-samples t-test). Summary statistics for each sequence and dataset during S1 are presented in Table S1.

### SUPPLEMENTARY NOTES: EXPLICIT MEMORY TASK

Given that TMR was shown to promote the emergence of explicit knowledge the next morning[1], we also set out to test whether this is true after a longer period. However, we found no difference between the free recall of the cued and uncued sequence ($z = -0.568$, $p = 0.570$, Wilcoxon signed-rank test), suggesting no TMR effect on the explicit knowledge of the sequence 20 days post-encoding (Fig.S1). Nevertheless, performance on both sequences differed from chance (cued: $z = -4.14$, $p < 0.001$; uncued: $z = -4.29$, $p < 0.001$; Wilcoxon signed-rank test), indicating that the participants learned both sequences explicitly over the course of the experiment.

### SUPPLEMENTARY NOTES: QUESTIONNAIRES

The EHI confirmed that all participants were right-handed, as the laterality quotient score (ranging between -100 and +100, where the negative values indicate left-handers and positive right-handers) was +100% for all but one subject who scored +75%. PSQI global scores (on a 21-points scale) ranged between 1 and 7 points, with a mean of 3.67 ($\pm$0.28), indicating, on average, a ‘good quality’ of sleep[2]. The median answer to the SQ (with 1 and 9 indicating the highest and lowest level of alertness, respectively) was 2 (IQR: 1) for all sessions, suggesting similar levels of alertness throughout the study.

Participants did report hearing experimental sounds during the night: On a 3-points scale, the median answer was 2 (IQR: 2), with 33% of the participants not hearing any sounds (answer 1), 20% of the participants being unsure (answer 2), and 47% of the participants hearing them clearly (answer 3). However, when asked about the number of sounds they had heard, the maximum number selected was 4 (reported by 13% of the participants), with the median answer of 2 (IQR: 2) sounds.

### SUPPLEMENTARY TABLES:

***Table S1. SRTT summary statistics.***

Mean reaction times (± SEM) (in ms) *for the BH, LH and RH trials of the cued and uncued sequence blocks (24 blocks per sequence) as well as random blocks (2 blocks with tones matching the cued sequence and 2 blocks with tones matching the uncued sequence) during Session 1. Average reaction times* (± SEM) for the last 4 blocks of each sequence are shown as well. *BH: both hands; LH: left hand; RH: right hand. n = 30.*

| Dataset | Cued sequence | Uncued sequence | Cued random | Uncued random | Cued sequence (last 4 blocks) | Uncued sequence (last 4 blocks) |
|---|---|---|---|---|---|---|
| **BH** | 321.82 ± 6.46 | 322.89 ± 7.83 | 357.81 ± 6.63 | 360.10 ± 6.69 | 286.05 ± 8.09 | 292.64 ± 9.99 |
| **LH** | 335.98 ± 7.18 | 334.90 ± 7.82 | 376.86 ± 7.45 | 371.60 ± 6.60 | 297.44 ± 9.19 | 303.92 ± 10.55 |
| **RH** | 307.67 ± 6.28 | 310.92 ± 8.44 | 347.77 ± 6.95 | 348.60 ± 7.76 | 274.62 ± 7.81 | 281.40 ± 10.20 |

***Table S2. Effect of and interaction between TMR, hand and session.***

*Results of the likelihood ratio tests between the full, linear mixed effects model and reduced models, i.e., models without the fixed effect of interest, or with an interaction. The full model was used to test the effect of TMR, hand and session on the early and late SSS. df: degrees of freedom; χ2: chi-squared; AIC: Akaike Information Criterion; SSS: Sequence Specific Skill. *p < 0.05.*

| | df | χ2 | p-value | AIC of a reduced model | AIC of a full model |
|---|---|---|---|---|---|
| **A. Both hands** | | | | | |
| i) Early SSS | | | | | |
| TMR | 1 | 1.5450 | 0.2138 | 1632.4 | 1632.9 |
| Session | 2 | 175.77 | **<0.0001*** | 1804.6 | 1632.9 |
| TMR x Session | 2 | 0.0740 | 0.9637 | 1636.3 | 1632.9 |
| ii) Late SSS | | | | | |
| TMR | 1 | 11.009 | **0.0009*** | 1621.3 | 1612.3 |
| Session | 2 | 93.041 | **<0.0001*** | 1701.3 | 1612.3 |
| TMR x Session | 2 | 3.0133 | 0.2216 | 1613.3 | 1612.3 |
| **B. Left hand** | | | | | |
| i) Early SSS | | | | | |
| TMR | 1 | 0.1878 | 0.6648 | 1666.5 | 1668.3 |
| Session | 2 | 132.02 | **<0.0001*** | 1796.3 | 1668.3 |
| TMR x Session | 2 | 0.1044 | 0.9492 | 1672.2 | 1668.3 |
| ii) Late SSS | | | | | |
| TMR | 1 | 4.015 | **0.0451*** | 1646.3 | 1644.2 |
| Session | 2 | 64.529 | **<0.0001*** | 1704.8 | 1644.2 |
| TMR x Session | 2 | 1.9750 | 0.3725 | 1646.3 | 1644.2 |
| **C. Right hand** | | | | | |
| i) Early SSS | | | | | |
| TMR | 1 | 3.2309 | 0.0723 | 1651.0 | 1659.8 |
| Session | 2 | 181.60 | **<0.0001*** | 1827.4 | 1649.8 |
| TMR x Session | 2 | 0.3741 | 0.8294 | 1653.4 | 1649.8 |
| ii) Late SSS | | | | | |
| TMR | 1 | 15.458 | **<0.0001*** | 1636.8 | 1650.3 |
| Session | 2 | 96.024 | **<0.0001*** | 1728.9 | 1636.2 |
| TMR x Session | 2 | 3.9605 | 0.1380 | 1636.9 | 1636.8 |
| **D. Left and right hand combined** | | | | | |
| i) Early SSS | | | | | |

| | | | | | |
|---|---|---|---|---|---|
| Hand | 1 | 22.423 | **<0.0001*** | 3286.3 | 3265.9 |
| Hand x Session | 2 | 9.1678 | **0.0102*** | 3260.7 | 3265.9 |
| TMR x Hand | 1 | 0.8955 | 0.3440 | 3267.0 | 3265.9 |
| ii) Late SSS | | | | | |
| Hand | 1 | 25.927 | **<0.0001*** | 3261.5 | 3237.5 |
| Hand x Session | 2 | 6.4262 | **0.0402*** | 3237.5 | 3235.9 |
| TMR x Hand | 1 | 2.1656 | 0.1411 | 3237.4 | 3237.5 |

***Table S3. Effect of session on SSS.***

*Post-hoc pairwise comparisons between sessions for the early and late SSS, conducted on each dataset separately. P-values reported are Holm adjusted. SSS: Sequence Specific Skill; S2-4: Session 2-4; df: degrees of freedom. *p < 0.05.*

| | **Mean S2 (±SE) [ms]** | **Mean S3 (±SE) [ms]** | **Mean S4 (±SE) [ms]** | **Estimate (±SE)** | **df** | **t ratio** | **p-value** (Holm adj) | **Effect size** |
|---|---|---|---|---|---|---|---|---|
| **A. Both hands** | | | | | | | | |
| i) Early SSS | | | | | | | | |
| S2-S3 | 39.2 (9.88) | 81.7 (10.20) | - | -42.5 (6.49) | 135 | -6.553 | **<0.0001*** | -1.320 |
| S3-S4 | - | 81.7 (10.20) | 163.1 (10.27) | -81.3 (6.63) | 131 | -12.270 | **<0.0001*** | -2.520 |
| ii) Late SSS | | | | | | | | |
| S2-S3 | 129.0 (8.92) | 154.0 (9.24) | - | -24.7 (6.14) | 135 | -4.018 | **0.0001*** | -0.807 |
| S3-S4 | - | 154.0 (9.24) | 200 (9.30) | -46.2 (6.27) | 131 | -7.367 | **<0.0001*** | -1.513 |
| **B. Left hand** | | | | | | | | |
| i) Early SSS | | | | | | | | |
| S2 – S3 | 35.3 (11.10) | 73.3 (11.40) | - | -37.9 (7.34) | 135 | -5.227 | **<0.0001*** | -1.050 |
| S3 – S4 | - | 73.3 (11.40) | 144.1 (11.5) | -70.8 (7.50) | 131 | -9.553 | **<0.0001*** | -1.960 |
| ii) Late SSS | | | | | | | | |
| S2 – S3 | 121.0 (10.20) | 149.0 (10.50) | - | -27.4 (6.74) | 135 | -4.064 | **0.0001*** | -0.817 |
| S3 – S4 | - | 149.0 (10.50) | 183 (10.6) | -34.4 (6.89) | 131 | -4.996 | **<0.0001*** | -1.026 |
| **C. Right hand** | | | | | | | | |
| i) Early SSS | | | | | | | | |
| S2 – S3 | 43.0 (9.13) | 90.0 (9.53) | - | -46.9 (7.13) | 136 | -6.627 | **<0.0001*** | -1.330 |
| S3 – S4 | - | 90.0 (9.53) | 182 (9.61) | -91.9 (7.30) | 132 | -12.670 | **<0.0001*** | -2.600 |
| ii) Late SSS | | | | | | | | |
| S2 – S3 | 137.0 (8.27) | 159.0 (8.69) | - | -21.9 (6.90) | 137 | -3.167 | **0.0019*** | -0.634 |
| S3 – S4 | - | 159.0 (8.69) | 217 (8.77) | -58.1 (7.08) | 132 | -8.207 | **<0.0001*** | -1.685 |

***Table S4. Effect of TMR on late SSS during each session.***

*Post-hoc pairwise comparisons of late SSS between the cued and uncued sequence on each session (S2-S4), conducted on each dataset separately. Both the uncorrected and Holm adjusted p-values are reported. S2-4: Session 2-4; SSS: Sequence Specific Skill; df: degrees of freedom. *p < 0.05. ^p < 0.07.*

| **Cued vs Uncued** | **Mean cued (± SE) [ms]** | **Mean uncued (± SE) [ms]** | **Estimate (± SE)** | **df** | **t ratio** | **p value** | **p value** (Holm adj) | **Effect size** |
|---|---|---|---|---|---|---|---|---|
| **A. Both hands** | | | | | | | | |
| S2 | 135 (9.94) | 123 (9.94) | -11.78 (7.95) | 133 | -1.482 | 0.1408 | 0.2815 | -0.390 |
| S3 | 157 (8.70) | 147 (8.70) | -9.98 (8.71 | 133 | -1.146 | 0.2537 | 0.2815 | -0.331 |
| S4 | 215 (11.60) | 185 (11.60) | -29.13 (8.89) | 133 | -3.277 | **0.0013*** | **0.0040*** | -0.965 |
| **B. Left hand** | | | | | | | | |
| S2 | 127 (11.20) | 116 (11.20) | -11.02 (8.76) | 133 | -1.258 | 0.2106 | 0.4212 | -0.3212 |
| S3 | 146 (10.30) | 145 (10.30) | -1.33 (9.60) | 133 | -0.139 | 0.8897 | 0.8897 | -0.0401 |
| S4 | 193 (13.10) | 172 (13.10) | -20.29 (9.79) | 133 | -2.072 | **0.0401*** | 0.1206 | -0.6101 |
| **C. Right hand** | | | | | | | | |
| S2 | 143 (9.73) | 130 (9.73) | -12.50 (8.94) | 133 | -1.401 | 0.1634 | 0.1634 | -0.369 |
| S3 | 167 (8.37) | 149 (8.37) | -18.60 (9.79) | 133 | -1.899 | 0.0597^ | 0.1193 | -0.548 |
| S4 | 237 (11.00) | 198 (11.00) | -38.20 (9.99) | 133 | -3.822 | **0.0002*** | **0.0006*** | -1.125 |

***Table S5. Effect of time and hand on the cueing benefit.***

*Results of the likelihood ratio tests between the full, linear mixed effects model and reduced models, i.e., models without the fixed effect of interest, or with an interaction. The full model was used to test the effect of hand and number of days post-TMR ('Time') on the cueing benefit (SSS for the uncued sequence subtracted from the cued sequence). χ2: chi-squared; AIC: Akaike Information Criterion. df: degrees of freedom. *p < 0.05. ^p < 0.07.*

| | df | χ2 | p-value | AIC of a reduced model | AIC of a full model |
|---|---|---|---|---|---|
| **A. Both hands** | | | | | |
| Time | 2 | 3.965 | **0.046*** | 809.14 | 807.18 |
| **B. Left hand** | | | | | |
| Time | 2 | 0.736 | 0.391 | 825.46 | 826.73 |
| **C. Right hand** | | | | | |
| Time | 2 | 6.581 | **0.010*** | 834.01 | 829.43 |
| **D. Left and right hand combined** | | | | | |
| Hand | 1 | 3.760 | 0.052^ | 1641.70 | 1639.90 |
| Hand x Time | 2 | 1.424 | 0.233 | 1639.9 | 1640.8 |

***Table S6. Sleep parameters.***

*Total recording duration, total sleep time, time spent in each sleep stage and time scored as movement presented as average (minutes ± SEM) and as percentage of the total recording duration. Total sleep time was calculated by subtracting the time spent awake from the total recording duration. N1-N3: stage 1 – stage 3 of NREM sleep. REM: Rapid Eye Movement sleep. n = 29.*

| | Percentage of total recording duration [%] | Mean duration ± SEM [min] |
|---|---|---|
| **Total recording duration** | 100% | 524.19 ± 10.29 |
| **Total sleep time** | 88.38% | 463.29 ± 12.89 |
| **Wake** | 11.50% | 60.90 ± 10.37 |
| **N1** | 4.52% | 23.33 ± 1.73 |
| **N2** | 46.35% | 242.45 ± 8.41 |
| **N3** | 19.91% | 104.22 ± 4.05 |
| **REM** | 15.86% | 83.43 ± 4.30 |
| **Movement** | 1.61% | 8.43 ± 1.32 |

***Table S7. Cueing benefit and the duration of N2 and N3.***

*Results of Pearson's correlations between cueing benefit and the percentage of time spent in N2 and N3. Both the uncorrected and FDR corrected p-values are reported. df: degrees of freedom; S2-4: Session 2-4; SSS: Sequence Specific Skill; N2-3: Stage 2-3 of NREM sleep. *p < 0.05.*

| | **Time spent in N2 [%]** | | | | **Time spent in N3 [%]** | | | |
|---|---|---|---|---|---|---|---|---|
| | df | Pearson's correlation | p-value | p-value (FDR corr) | df | Pearson's correlation | p-value | p-value (FDR corr) |
| **A. Both hands** | | | | | | | | |
| S2 | 25 | 0.197 | 0.324 | 0.324 | 27 | -0.015 | 0.939 | 0.939 |
| S3 | 20 | 0.378 | 0.082 | 0.144 | 21 | -0.031 | 0.887 | 0.939 |
| S4 | 19 | 0.372 | 0.096 | 0.144 | 21 | 0.089 | 0.697 | 0.939 |
| **B. Left hand** | | | | | | | | |

| | | | | | | | | |
|---|---|---|---|---|---|---|---|---|
| S2 | 23 | 0.101 | 0.632 | 0.632 | 25 | 0.036 | 0.857 | 0.974 |
| S3 | 19 | 0.432 | **0.045*** | 0.135 | 21 | 0.070 | 0.746 | 0.974 |
| S4 | 19 | 0.324 | 0.152 | 0.228 | 21 | 0.007 | 0.974 | 0.974 |
| **C. Right hand** | | | | | | | | |
| S2 | 24 | -0.030 | 0.884 | 0.884 | 26 | -0.053 | 0.788 | 0.788 |
| S3 | 19 | 0.177 | 0.431 | 0.657 | 21 | -0.125 | 0.562 | 0.788 |
| S4 | 19 | 0.377 | 0.092 | 0.276 | 21 | 0.144 | 0.511 | 0.788 |

***Table S8. Summary statistics for sleep spindles.***

*Average number and density (number/min) of spindles (± SEM). Results are presented for N2 and N3 of the cue (A) and no-cue (B) period. N2-3: stage 2-3 of NREM sleep.*

| | **Density** | | | **Number** | | |
|---|---|---|---|---|---|---|
| | Average | Left | Right | Average | Left | Right |
| **A. Cue** | | | | | | |
| N2 | 5.28 ±0.27 | 5.43 ±0.29 | 5.13 ±0.19 | 124.02 ±11.31 | 127.68 ±11.70 | 120.35 ±11.23 |
| N3 | 4.24 ±0.16 | 4.31 ±0.16 | 4.17 ±0.16 | 191.58 ±13.14 | 195.39 ±13.88 | 187.77 ±12.74 |
| N2 & N3 | 4.20 ±0.22 | 4.43 ±0.25 | 3.96 ±0.24 | 286.91 ±18.50 | 301.06 ±19.81 | 272.76 ±18.70 |
| **B. No-Cue** | | | | | | |
| N2 | 4.85 ±0.27 | 5.02 ±0.29 | 4.68 ±0.28 | 51.71 ±5.07 | 53.17 ±5.22 | 50.24 ±5.08 |
| N3 | 3.86 ±0.16 | 3.94 ±0.17 | 3.78 ±0.17 | 79.48 ±5.68 | 81.30 ±5.79 | 77.66 ±5.69 |
| N2 & N3 | 3.80 ±0.20 | 4.01 ±0.23 | 3.59 ±0.21 | 117.74 ±7.69 | 123.59 ±8.12 | 111.89 ±7.79 |

***Table S9. Cueing benefit and spindle density.***

*Results of Pearson's (A) and Spearman's (B) correlations (both FDR corrected and not) between the cueing benefit for BH dataset during each of the post-stimulation sessions (S2, S3, S4) and spindle density averaged over all motor channels during the cue (A) and no-cue (B) period. N2 (green) and N3 (blue) were analysed separately and together (N23, purple). df: degrees of freedom; S2-4: Session 2-4; N2-N3: stage 2 - stage 3 of NREM sleep; BH: Both Hands.*

| **Dataset** | **Session** | **Sleep stage** | **Correlation coefficient** | **p-value** | **df** | **p-value (FDR corr)** |
|---|---|---|---|---|---|---|
| | | | **A. Cue period** | | | |
| BH | S2 | N2 | 0.237 | 0.224 | 26 | 0.288 |
| BH | S3 | N2 | -0.254 | 0.231 | 22 | 0.288 |
| BH | S4 | N2 | -0.231 | 0.288 | 21 | 0.288 |
| BH | S2 | N3 | -0.04 | 0.840 | 25 | 0.840 |
| BH | S3 | N3 | -0.337 | 0.126 | 20 | 0.192 |
| BH | S4 | N3 | -0.343 | 0.128 | 19 | 0.192 |
| BH | S2 | N23 | 0.196 | 0.309 | 27 | 0.322 |
| BH | S3 | N23 | -0.277 | 0.191 | 22 | 0.322 |
| BH | S4 | N23 | -0.216 | 0.322 | 21 | 0.322 |
| | | | **B. No-cue period** | | | |
| BH | S2 | N2 | 0.135 | 0.485 | 27 | 0.485 |
| BH | S3 | N2 | -0.317 | 0.132 | 22 | 0.396 |
| BH | S4 | N2 | -0.227 | 0.296 | 21 | 0.444 |
| BH | S2 | N3 | 0.227 | 0.235 | 27 | 0.546 |
| BH | S3 | N3 | -0.171 | 0.422 | 22 | 0.546 |
| BH | S4 | N3 | -0.132 | 0.546 | 21 | 0.546 |

| | | | | | | |
|---|---|---|---|---|---|---|
| BH | S2 | N23 | 0.189 | 0.326 | 27 | 0.639 |
| BH | S3 | N23 | -0.172 | 0.419 | 22 | 0.629 |
| BH | S4 | N23 | -0.042 | 0.851 | 21 | 0.851 |

***Table S10. Functional TMR-related activity.***

*Clusters showing increased (inc) or decreased (dec) activity for the cued relative to uncued sequence alone (A - S2 and S3, B - S2) or when considering covariates of cueing benefit during S2 (C) and cueing benefit during S4 (E). (D) Decrease in activity from S2 to S3 for the [cued > uncued] contrast when considering behavioural cueing benefit at S4 as a covariate. No significant clusters were found when considering cueing benefit during S3 as a covariate.*

| Region | MNI x, y, z (mm) | Number of voxels | F/T peak | $P_{FWE}$ peak |
|---|---|---|---|---|
| **A.** Main effect of TMR for S2 and S3 | | | | |
| i. Right Precuneus (inc) | 8, -72, 58 | 9 | 22.67 | 0.032 |
| **B.** [Cued > Uncued at S2] | | | | |
| i. Left Precuneus (inc) | -9 -62, 66 | 9 | 4.79 | 0.020 |
| **C.** [Cued > Uncued at S2 * cueing benefit at S2] | | | | |
| i. Left Precuneus (inc) | -4, -78, 46 | 40 | 5.18 | 0.009 |
| ii. Left Precuneus (inc) | -18, -68, 36 | 1 | 4.44 | 0.046 |
| iii. Right Cerebellum (inc) | 28, -58, -30 | 1 | 4.94 | 0.044 |
| iv. Left Putamen (inc) | -24, 4, 6 | 3 | 4.41 | 0.034 |
| **D.** [Δ Cued > Uncued from S2 to S3 * cueing benefit at S4] | | | | |
| i. Left Precuneus (dec) | -2, -54, 10 | 18 | 6.66 | 0.002 |
| **E.** [Cued > Uncued at S3 * cueing benefit at S4] | | | | |
| i. Right Postcentral gyrus (inc) | 58, -18, 38 | 7 | 5.50 | 0.022 |
| ii. Left Parahippocampus (dec) | -22, -26, -16 | 1 | 4.52 | 0.047 |

*Regions listed were significant at peak voxel threshold of $p_{FWE} < 0.05$, after correction for multiple voxel-wise comparisons within pre-defined bilateral ROI for bilateral precuneus (A.i, B.i, C.i, C.ii, D.i), bilateral hippocampus and parahippocampus (E.ii), bilateral cerebellum (C.iii), bilateral dorsal striatum (C.iv), bilateral sensorimotor cortex (E.i). Peak voxel MNI coordinates and peak F (A) and T (B-E) values are given. n = 22 for (A), n = 28 for (B) and (C), n = 23 for (E), n = 21 for (D).*

***Table S11. Structural brain changes over time.***

*Clusters showing increased (inc) or decreased (dec) changes in grey matter volume over time and when considering covariates of cueing benefit during S3 and S4. No significant clusters were found when considering cueing benefit during S2 as a covariate. No significant clusters were found in white matter either.*

| Region | MNI x, y, z (mm) | Number of voxels | T peak | $P_{FWE}$ peak |
|---|---|---|---|---|
| A. [Δ GM volume from S1 to S3 * cueing benefit at S4] | | | | |
| Right Precentral gyrus (inc) | 42, -2, 45 | 5 | 6.21 | 0.020 |
| B. [Δ GM volume from S1 to S3 * cueing benefit at S3] | | | | |
| Left Fusiform (dec) | -28, - 26, -24 | 5 | 5.51 | 0.025 |

*Regions listed were significant at peak voxel threshold of $p_{FWE} < 0.05$, after correction for multiple voxel-wise comparisons within pre-defined ROI for bilateral sensorimotor cortex (A) and bilateral hippocampus and parahippocampus (B). Since fusiform gyrus was not our ROI, the result in (B) has likely arisen due to the imperfection of the method and is thus not discussed any further. Peak voxel MNI coordinates, and peak T values are given. n = 24 for (A), n = 29 for (B).*

### <u>SUPPLEMENTARY FIGURES:</u>

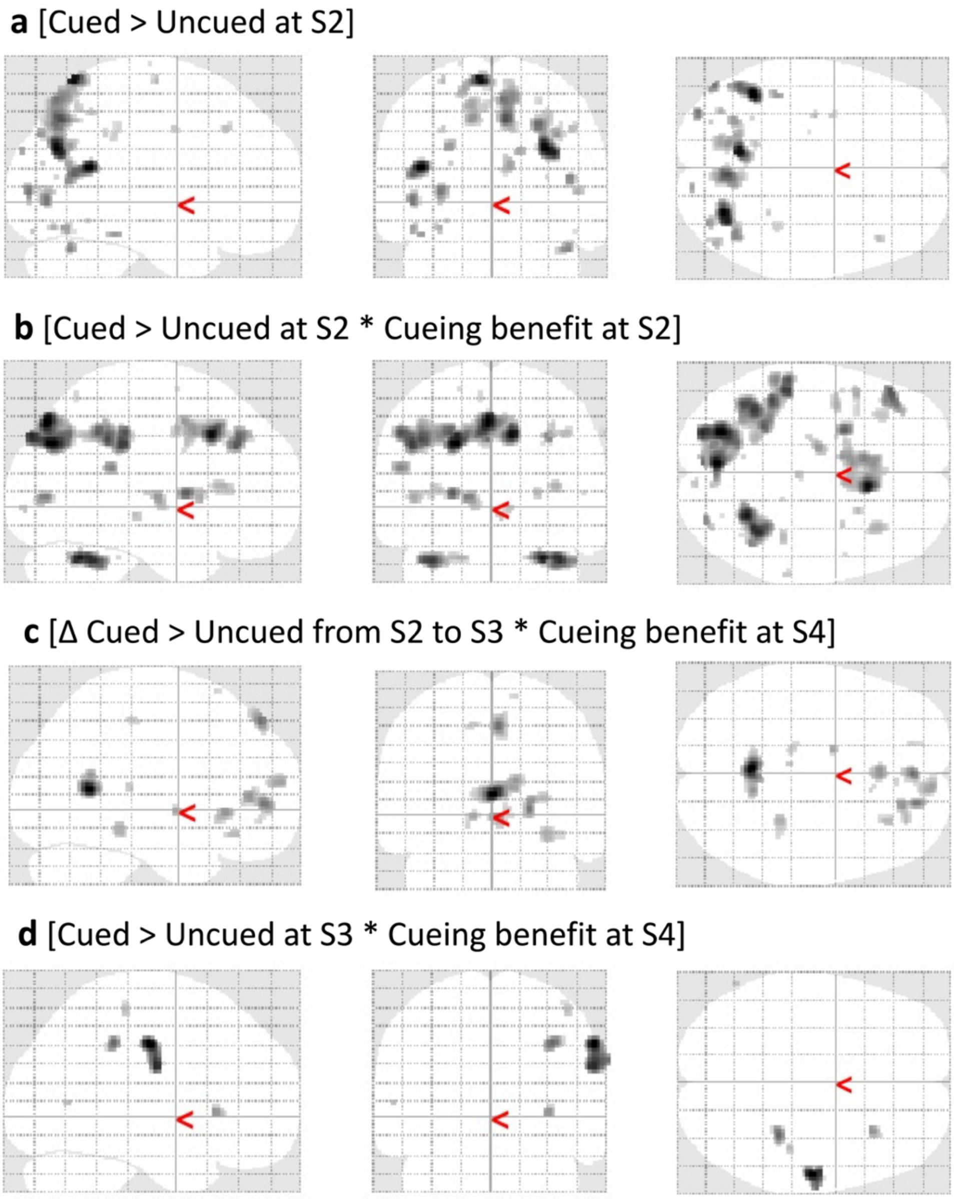


***Fig. S1. Glass brain fMRI results.** SPM fMRI results in glass brain projection displayed at $p < 0.001$, uncorrected, for the same contrasts as in Fig.5 and Fig.6A-B. **(A)** TMR-dependent increase in brain activity 24 h post-stimulation. **(B)** Brain activity for the cued > uncued contrast at S2 was positively associated with behavioural cueing benefit at the same time point. **(C)** A change in brain activity from S2 to S3 for the cued > uncued contrast was negatively associated with behavioural cueing benefit at S4. **(D)** Brain activity for the cued > uncued contrast at S3 was positively associated with behavioural cueing benefit at S4. S2-4: Session 2-4; n = 28 for (A-B), n = 21 for (C), n = 23 for (D).*

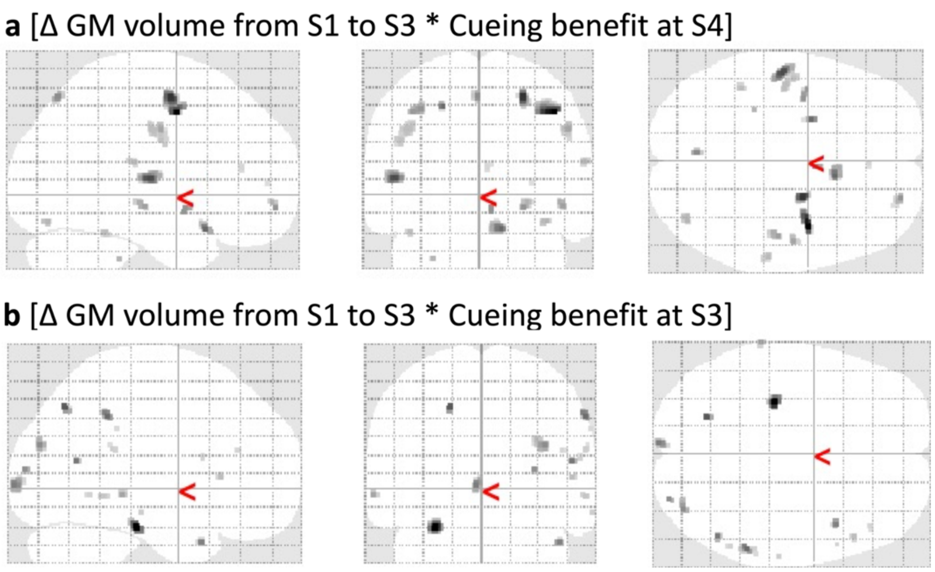


***Fig. S2. Glass brain VBM results.*** *SPM VBM results in glass brain projection displayed at $p < 0.001$, uncorrected, for the same contrasts as in Fig.6C-D (A) and Table S11 (A, B).* ***(A)*** *An increase in grey matter volume at S3 relative to S1 was associated with an increase in behavioural cueing benefit at S4.* ***(B)*** *Reduction in grey matter volume at S3 relative to S1 associated with cueing benefit at S3. S1-4: Session 1-4. n = 24 for (A), n = 29 for (B).*

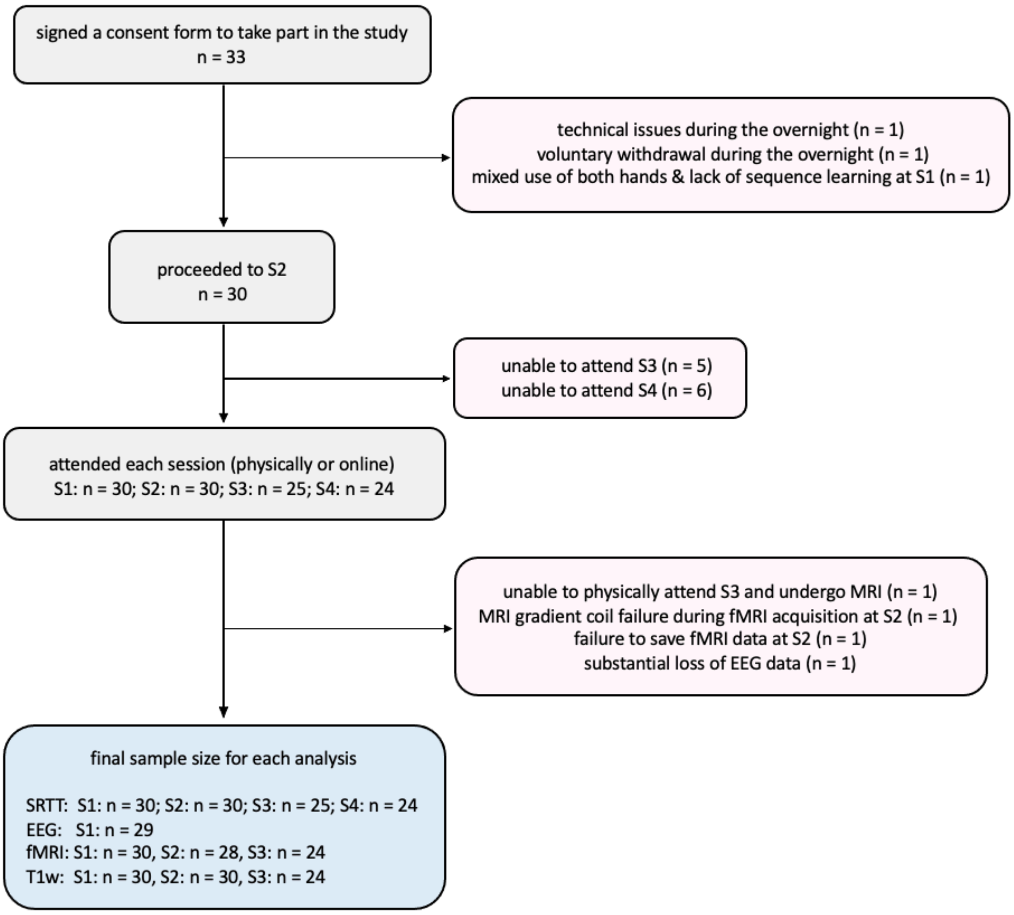


***Fig.S3. A flowchart illustrating participants included and excluded from the analysis.*** *On the left, the number of participants included in the study at different time points, with the final sample size shown at the bottom. On the right, the number of participants excluded (in red) together with a reason for the exclusion.*

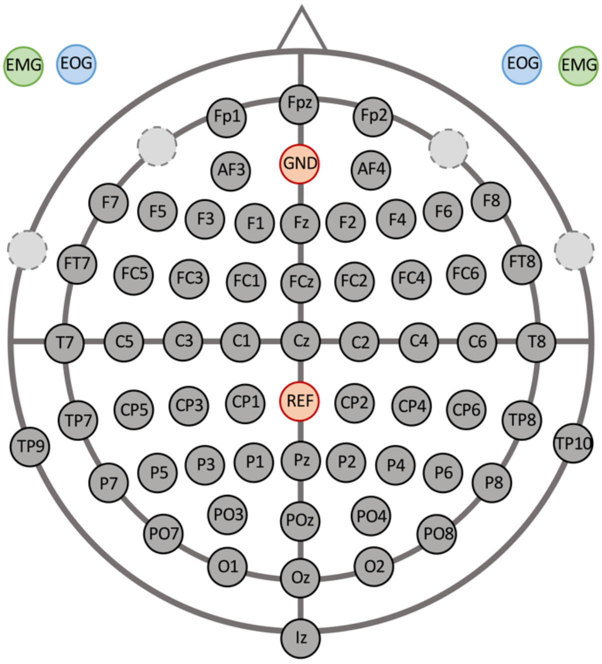


***Fig. S4. EEG electrodes layout.*** *Orange: ground (GND) and reference (REF) electrodes; light grey (dashed circles): original position of the electrodes used to record EMG and EOG; green: EMG electrodes; blue: EOG electrodes.*

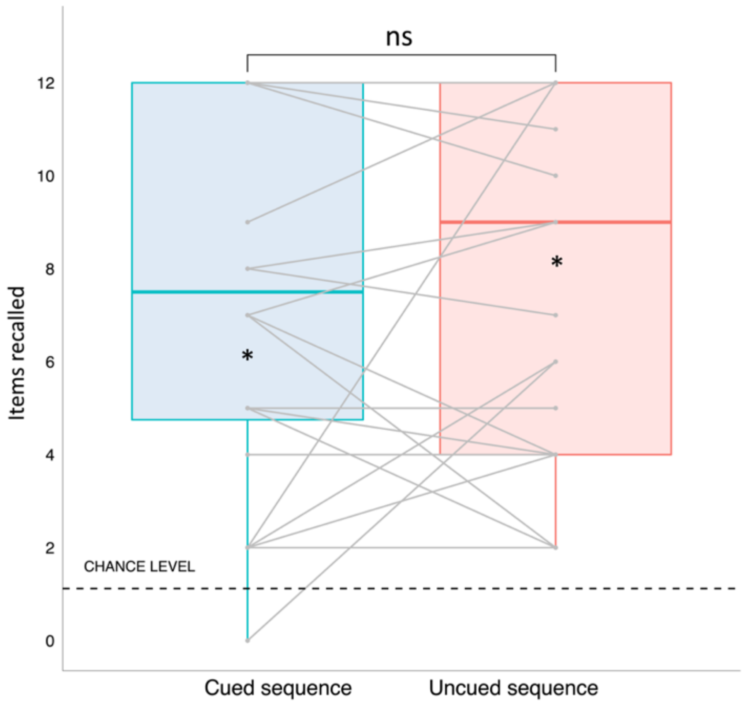


***Fig. S5. Cueing memory reactivation during sleep does not affect explicit memory of the sequence.*** *Explicit knowledge of both sequences was significantly above chance (significance denoted with *) 20 days post-encoding, although no effect of TMR was evident. Geoms represent median ±IQR for the cued (blue) and uncued (red) sequence, whiskers represent largest and lowest values within 1.5 IQR above and below the 75$^{th}$ and the 25$^{th}$ percentile, respectively. Grey dots represent performance of each subject. ns: non-significant. n = 24.*
